# Structural insights on the substrate-binding proteins of the *Mycobacterium tuberculosis* mammalian-cell-entry (Mce) 1 and 4 complexes

**DOI:** 10.1101/2020.09.29.317909

**Authors:** Pooja Asthana, Dhirendra Singh, Jan Skov Pedersen, Mikko J. Hynönen, Ramita Sulu, Abhinandan V. Murthy, Mikko Laitaoja, Janne Jänis, Lee W. Riley, Rajaram Venkatesan

**Affiliations:** Faculty of Biochemistry and Molecular Medicine, University of Oulu, Oulu, Finland; Department of Chemistry and Interdisciplinary Nanoscience Center (iNANO), Aarhus University, Aarhus, Denmark; Faculty of Medicine, University of Helsinki, Finland; Department of Chemistry, University of Eastern Finland, Joensuu, Finland; School of Public Health, University of California, Berkeley, CA, USA

**Keywords:** *Mycobacterium tuberculosis*, mammalian-cell-entry proteins, Mce1, Mce4, substrate-binding proteins, ABC/lipid transporter, crystal structure, SAXS

## Abstract

Tuberculosis (Tb), caused by *Mycobacterium tuberculosis* (*Mtb*), is responsible for more than a million deaths annually. In the latent phase of infection, *Mtb* uses lipids as the source of carbon and energy for its survival. The lipid molecules are transported across the cell wall via multiple transport systems. One such set of widely present and less-studied transporters is the Mammalian-cell-entry (Mce) complexes. Here, we report the properties of the substrate-binding proteins (SBPs; MceA-F) of the Mce1 and Mce4 complexes from *Mtb* which are responsible for the import of mycolic acid/fatty acids, and cholesterol respectively. MceA-F are composed of four domains namely, transmembrane, MCE, helical and tail domains. Our studies show that MceA-F are predominantly monomeric when purified individually and do not form homohexamers unlike the reported homologs (MlaD, PqiB and LetB) from other prokaryotes. The crystal structure of MCE domain of *Mtb* Mce4A (MtMce4A_39-140_) determined at 2.9 Å shows the formation of an unexpected domain-swapped dimer in the crystals. Further, the purification and small-angle X-ray scattering (SAXS) analysis on MtMce1A, MtMce4A and their domains suggest that the helical domain requires hydrophobic interactions with the detergent molecules for its stability. Combining all the experimental data, we propose a heterohexameric arrangement of MtMceA-F SBPs, where the soluble MCE domain of the SBPs would remain in the periplasm with the helical domain extending to the lipid layer forming a hollow channel for the transport of lipids across the membranes. The tail domain would reach the cell surface assisting in lipid recognition and binding.

## 1. Introduction

*Mycobacterium tuberculosis* (*Mtb*) is a deadly intracellular pathogen causing the disease tuberculosis (Tb) and is responsible for more than a million deaths every year. Approximately, one fourth of the world population is latently infected by *Mtb* (1). *Mtb* is one of the very few bacteria which can use host lipids as the source of energy and carbon, and this property might be most relevant during the intra-phagosomal latent stage of infection (2). Interestingly, *Mtb* has nearly 200 lipid metabolizing proteins (3, 4). Therefore, lipid transport across the membrane plays a pivotal role in *Mtb* pathogenesis. One such set of lipid transporters found in *Mtb* is encoded by the, <u>M</u>ammalian-<u>c</u>ell-<u>e</u>ntry (Mce) operons, Mce1, Mce2, Mce3, and Mce4 (4), comprising 10-14 genes each. These operons were named based on the initial observation that a DNA fragment (corresponding to Mce1A) from H37Ra when expressed in *E.coli* made them to invade HeLa cells (5). Similar to Mce1A, the expression of Mce3A and Mce4A in *E. coli* also provides the invasion ability to enter HeLa cells (6, 7). However, further studies have suggested that Mce operons encode for substrate-binding proteins (SBPs) (MceA-F) and permeases (YrbEA-B) to form an ABC transporter (8, 9). The ATPase for this ABC-transporter is proposed to be coded by the gene MceG (also known as Mkl) elsewhere (10). In addition, Mce1, Mce3 and Mce4 operons code for Mce-associated membrane proteins (Mam, also known as Mas) which probably stabilizes the Mce complexes (8). It is now well demonstrated that Mce1 is involved in the transport of mycolic acid/fatty acid and Mce4 imports cholesterol. While *Mtb* disrupted in Mce2 operon accumulates sulfolipid-1 at levels nearly 10 times that of wild type *Mtb* during stationary growth (2, 9, 11), the substrate specificity for Mce3 complex is still unknown. The role of protein encoded by Mce operons in the pathogenicity of *Mtb* has been well established in mouse models based on operon mutants (12, 13).

Recently, structures of homologs of Mce SBPs from *E.coli* (MlaD, PqiB, LetB) and *A. baumannii* (MlaD) were determined (14–16). Based on these homohexameric structures, two different mechanism of lipid transport has been reported. First, Mla complex-ferry transport mechanism, where the Mla operon carries a single Mce gene (MlaD) with a single MCE domain. In this case, the lipids are transferred to MlaD by a shuttle protein (MlaC) and the lipid molecule is then passed through the central hydrophobic pore of homohexameric MlaD. Second, the LetB and PqiB tunnel transport mechanism, where LetB forms a long stack of seven homohexameric MCE domains one above the other connecting the inner and outer membrane with central channel mediating lipid transport. Like LetB, PqiB also forms a central pore formed by three stacked Mce homohexamers with their long C-terminal helix forming a narrow channel for lipid transport.

In this study, we have recombinantly expressed and purified the SBPs MtMce1A-1F as well as MtMce4A-4F encoded in the Mce1 and Mce4 operons. In addition, based on sequence analysis and secondary structure prediction, various domains of MtMce1A and MtMce4A have been identified and a detailed characterization including small-angle X-ray scattering (SAXS) analysis has been performed. Furthermore, the crystal structure of the MCE domain of MtMce4A has been determined using Selenomethionine-single-wavelength anomalous dispersion (SeMet-SAD) phasing which forms an unexpected domain-swapped dimer in the crystals. This is the first reported structure of any mycobacterial protein encoded by the Mce operons. In addition, our studies show that these proteins are predominantly monomeric when expressed and purified individually. Moreover, the purification and SAXS studies of MtMce1A and MtMce4A domains further suggests that the helical domain is mainly responsible for the interaction with the detergent micelles, which are required in the purification indicating that this region may form the hydrophobic pore for the transfer of lipid molecules in the complex. Based on these results, a possible model for the *Mtb* Mce complexes is provided.

## 2. Results

### 2.1 Mtb MceA-F SBPs have conserved four-domain architecture

A comparative sequence analysis and secondary structure prediction (*SI Appendix*, Fig. S1, S2) of *Mtb* Mce 1, 2, 3 and 4 A-F SBPs suggest that in spite of having very low sequence identity between them (~20% or less; *SI Appendix*, Table S1, S2), they have conserved domain architecture with each of them having four domains (Fig. 1*A* and 1*B*). The first domain is the N-terminal transmembrane (TM) domain (~30-40 amino acids) predicted to form a single transmembrane helix followed by the second domain with ~100 amino acids composed of mainly β-strands (usually 7 β-strands), now widely known as MCE domain. The third domain mainly consists of long helices (~200 amino acids) and is therefore referred here as ‘helical domain’. The fourth domain is mainly predicted to be an unstructured domain and referred as ‘tail domain’. Interestingly, the length of the tail domain varies between 6-260 amino acids between the various MtMceA-F SBPs, while the other domains are well conserved with respect to their topology and number of amino acids. For example, among the MtMce1A-1F proteins, the MtMce1E tail domain has 34 residues and MtMce1F has 218 residues. Additionally, the tail domains of MtMce1C, MtMce1D, MtMce4D and MtMce4F are proline-rich. Moreover, MtMce1E, MtMce2E, MtMce3E and MtMce4E act as probable lipoproteins also known as Lprk, LprL, LprM, and LprN respectively with a conserved ‘lipobox’ at the N-terminus (17).

**Fig.1:**
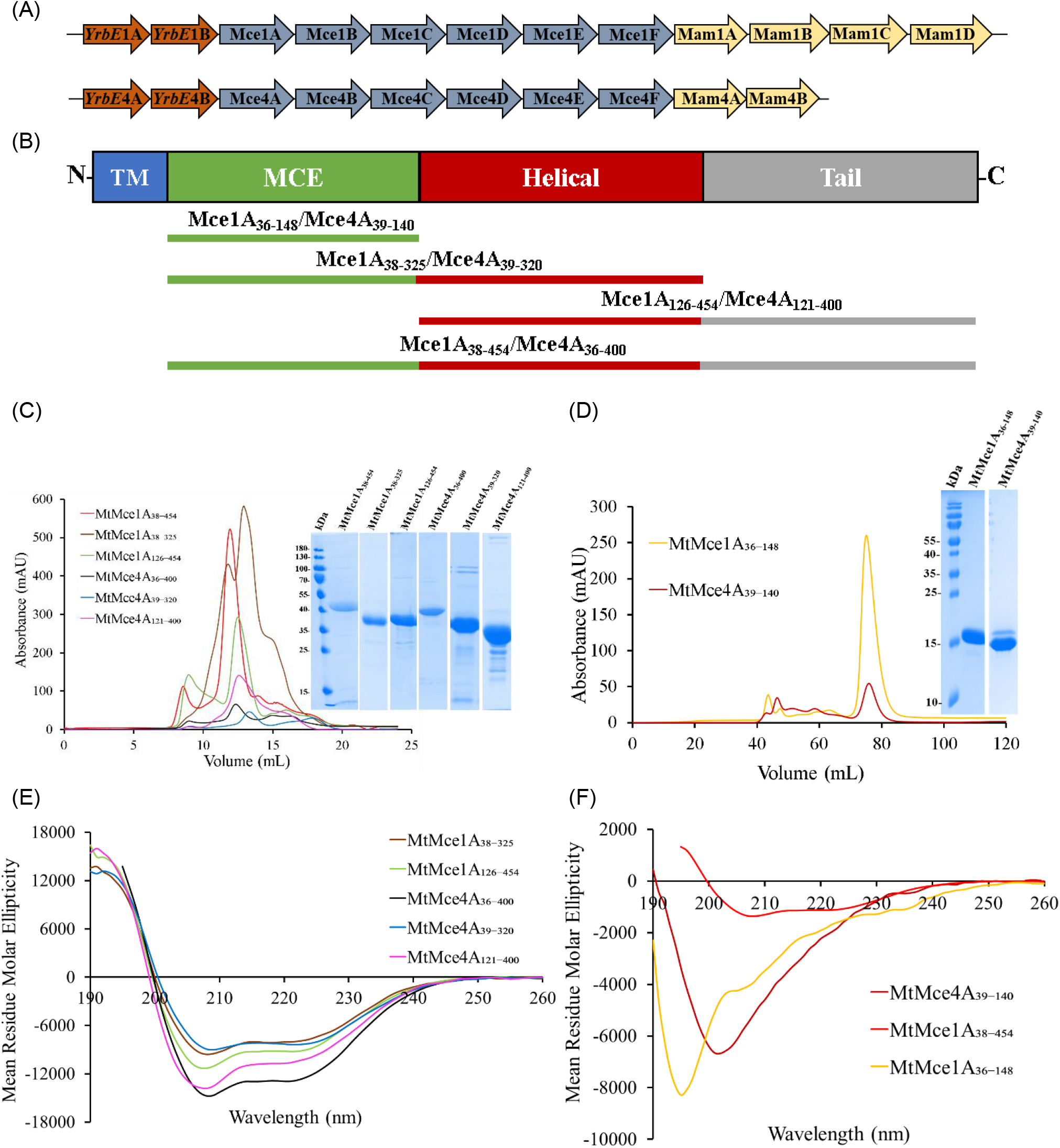
**(A)** Part of Mce1 and Mce4 operons of *Mtb* comprising permeases (YrbEA-B), SBPs (MceA-F) and Mam proteins. **(B)** Domain architecture of the individual Mce SBPs. The Mce SBPs are characterized by four domains named as transmembrane (TM), MCE, helical and tail domain. The constructs of MtMce1A and MtMce4A used in this study are mentioned below in the same color coding. **(C)** Size exclusion chromatography (SEC) elution profiles of MtMce1A_38-325_, MtMce1A_126-454_, MtMce1A_38-454_ and MtMce4A_39-320_, MtMce4A_121-400_, and MtMce4A_36-400_ on a 24 ml Superdex 200 10/300 column. The protein samples were analyzed on a 12 % SDS-PAGE (inset). **(D)** SEC elution profile of MtMce1A_36-148_ and MtMce4A_39-140_ on a 120 ml Superdex 75 HiLoad 16/600 column. The protein samples were analyzed on a 18% SDS-PAGE (inset) **(E)** The CD spectra of MtMce1A_38-325_ (brown), MtMce1A_126-454_ (green), MtMce4A_36-400_ (black), MtMce4A_39-320_ (blue) and MtMce4A_121-400_ (pink). **(F)** The CD spectra of MtMce1A_38-454_ (red), MtMce1A_36-148_ (yellow) and MtMce4A_39-140_ (maroon).

### 2.2 MCE domain is the only soluble part of MtMce1A and MtMce4A

All the six SBPs encoded in the Mce1 and Mce4 operons (MtMce1A-1F and MtMce4A-4F) were recombinantly expressed in *E. coli* and purified (*SI Appendix*, Supplementary Results, Fig. S3) in the presence of detergents. Given that, all the substrate-binding proteins have been predicted to have similar secondary structures and domain architecture, further detailed domain-level characterization was performed for MtMce1A and MtMce4A. From secondary structure predictions (Section 3.1), the domain constructs of MtMce1A and MtMce4A categorized as MCE (MtMce1A_36-148_, MtMce4A_39-140_), MCE+Helical (MtMce1A_38-325_, MtMce4A_39-320_), Helical+Tail (MtMce1A_126-454_, MtMce4A_121-400_), and MCE+Helical+Tail domains (MtMce1A_38-454_, MtMce4A_36-400_) were successfully expressed in *E. coli*, and screened to evaluate their solubility in the presence and absence of detergents. Interestingly, the MCE domain of both MtMce1A and MtMce4A (MtMce1A_36-148_ and MtMce4A_39-140_) were the only soluble constructs. Whereas, the MCE+Helical+Tail, MCE+Helical as well as Helical+Tail domains could be purified only in the presence of detergent even though the transmembrane domain has been deleted in all these constructs (Fig. 1*B* and *C*). Additionally, the extension of the soluble construct even with one (MtMce4A_39-154_) or two helical domains (MtMce4A_39-190_) resulted in insolubility, indicating that the helical domain requires detergent for its stability. The CD curves of MtMce1A and MtMce4A domains (Fig. 1*D* and *E*) indicated mixtures of α-helical and β-sheet content for all the MtMce1A and MtMce4A domains purified with DDM. Whereas, the soluble constructs (MtMce1A_36-148_ and MtMce4A_39-140_) showed a typical β-sheet dominated spectra (*SI Appendix*, Table S3).

### 2.3 MtMce4A_39-140_ crystallizes as a domain-swapped dimer

Structural studies were initiated on MtMce1A_38-454_, MtMce4A_36-400_ as well as soluble MCE domains MtMce1A_38-148_ and MtMce4A_39-140_. Despite extensive trials, only the MtMce4A_39-140_ crystallized readily in several conditions in the space group P65. Given the low sequence identity of MtMce4A_39-140_ with its homologous proteins (~15%), the structure of MtMce4A_39-140_ was determined using SeMet-SAD to 2.9 Å resolution. Although the initial Mathew’s coefficient calculations suggested the presence of 6-8 molecules in the asymmetric unit with a solvent content of about 43-57%, the solved structure showed that only four molecules are present in the asymmetric unit corresponding to a solvent content of about 71%. Interestingly, further refinement and model building of the structure revealed that the tetramer in the asymmetric unit is composed of two domain-swapped dimers (Fig. 2*A*). The domain-swapped dimer is formed by the extension of residues 107-141 from one molecule into the other molecule. The swapped region has two β-strands and an extended loop (Fig. 2*A*).

**Fig.2:**
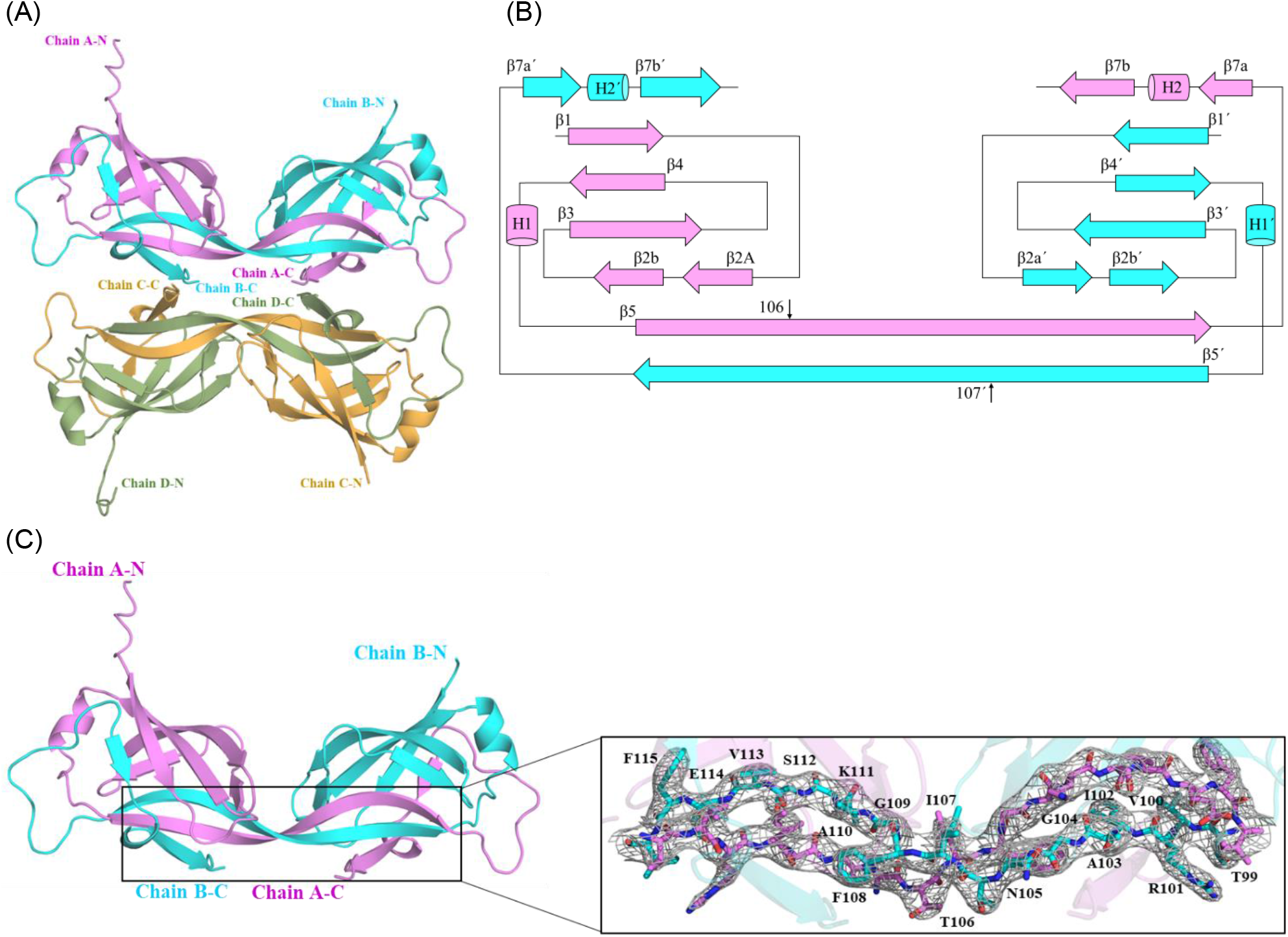
**(A)** Crystal structure of the MtMce4A_39-140_ with 4 molecules in the asymmetric unit. **(B)** Topology of MtMce4A_39-140_ domain-swapped dimer. The β-strands are shown as *arrows* and the helices as *cylinder*. The secondary structures of chain A and chain B are shown in *pink* and *cyan*, respectively. The secondary structure elements and residue numbers for chain B are indicated with prime (‘). The residues after the black vertical arrow are involved in the domain-swapping. **(C)** The domain-swapped dimer residues of β-5 and β-5′ are highlighted and shown in the inset. The 2Fo-Fc electron density maps contoured at 1.5 σ are shown in grey mesh. These residues are important for the arrangement of the domain-swapped dimer.

The topology diagram for the swapped dimer is shown in Fig. 2*B*. The residues involved in the formation of β-barrel are Thr40-Ser46 (β1), Leu52-Met54 (β2a), Lys59-Gly65 (β2b), Ile65-Ser74 (β3), Arg81-Asp87 (β4), Thr99-Thr106 (β5), Ile107-Ile116 (β5′) (considered as 6^th^ β-strand), His131-Val132 (β7a′), Val137-Glu141 (β7b′). The residues from 107-141 are exchanged between the two monomers for completing the signature MCE-fold. The overall structure has visible electron density for all the residues corresponding to MtMce4A_39-140_ except for the N-terminal residues 1-31, and C-terminal residues 143-146 corresponding to the vector region. The data collection and refinement statistics are mentioned in Table 1.

**Table 1:** Crystallization, data collection and refinement statistics of MtMce4A_39-140_ structure.

| Dataset | SeMet-labelled | Native |
| --- | --- | --- |
| **Crystallization** |  |  |
| Protein storage buffer | 50 mM MOPS, 350 mM NaCl, 10% Glycerol, pH 7.0 | 50 mM MOPS, 350 mM NaCl, 10% Glycerol, 5 mM DTT, pH 7.0 |
| Protein concentration | 7.5 mg ml<sup>-1</sup> | 7.5 mg ml<sup>-1</sup> |
| Crystallization conditions | 100 mM MES, 700 mM ammonium sulfate, pH 6.0 | 100 mM Sodium HEPES, 100 mM LiCl<sub>2</sub>, 20% PEG 400, pH 7.5 |
| Cryo-protectant | 25% Glycerol | 20% Ethylene Glycol |
| Temperature | 22 °C | 22 °C |
| **Data Collection:** |  |  |
| Beam line | BioMax | BioMax |
| Wavelength (Å) | 0.968 | 0.953 |
| Detector | Eiger 16M Hybrid-pixel | Eiger 16M Hybrid-pixel |
| Detector distance (mm) | 357.46 | 276.71 |
| Oscillation range (°) | 0.1 | 0.1 |
| **Data Processing:** |  |  |
| Space group | P6<sub>5</sub> | P6<sub>5</sub> |
| a, b, c (Å) | 134.0, 134.0, 105.5 | 131.2, 131.2, 105.5 |
| $\alpha$, $\beta$, $\gamma$ (°) | 90, 90, 120 | 90, 90, 120 |
| Resolution range (Å) | 48-3.6(3.9-3.6)* | 47.8-2.9 (3.0-2.9) * |
| R<sub>pim</sub> (%) | 0.11 (1.45) | 0.06 (0.79) |
| Multiplicity | 12.4 (12.5) | 15.4 (15.6) |
| Wilson B-factor (Å<sup>2</sup>) | 108.4 | 66.3 |
| Solvent content | 72.90 | 71.74 |
| Total number of reflections | 155214 (36878) | 355222 (57858) |
| Number of unique reflections | 12505 (2961) | 23016 (3714) |
| CC (1/2) | 99.6 (23.0) | 99 (43.6) |
| $\langle I \rangle / \sigma \langle I \rangle$ | 7.1 (1.2) | 11 (1.4) |
| Completeness (%) | 99.8 (99.5) | 100 (99.9) |
| **Refinement statistics** |  |  |
| R-work | 0.2165 | 0.1947 |
| R-free | 0.2466 | 0.2348 |
| Number of atoms |  |  |
| Protein | 3271 | 3276 |
| Water | - | 14 |
| Average B-factor (Å<sup>2</sup>) |  |  |
| Protein | 158.89 | 92.22 |
| Water | - | 79.08 |
| Ramachandran statistics |  |  |
| Allowed (%) | 6 | 3 |
| Favored (%) | 93.8 | 97 |
| Outliers (%) | 0.2 | 0 |
| R.M.S deviations |  |  |
| Bond lengths (Å) | 0.002 | 0.002 |
| Bond angles (°) | 0.560 | 0.422 |
| PDB id | 7AI2 | 7AI3 |

\*Values in parenthesis are for the highest resolution shell.

### 2.4 MtMce1A and MtMce4A are predominantly monomeric in solution

The domain-swapped dimeric nature of MtMce4A_39-140_ was unexpected. To further verify their oligomeric states in solution, SEC-multi angle light scattering (SEC-MALS) studies were conducted for all the purified MtMce1A and MtMce4A domains. Interestingly, these studies showed that MtMce1A and MtMce4A domains were predominantly monomeric in nature (Table 2 & *SI Appendix*, Fig. S4, S5). The MtMce1A and MtMce4A domains purified in DDM showed two peaks in the elution profile corresponding to protein-detergent complex (PDC) and empty detergent micelle. Whereas, MtMce1A_36-148_ and MtMce4A_39-140_ which are soluble and purified without DDM has a single scattering peak, corresponding to the monomeric molecular mass (*SI Appendix*, Fig. S4, S5). As the SEC-MALS analysis showed that both MtMce1A_36-148_ and MtMce4A_39-140_ are monomeric in solution and MtMce4A_39-140_ is a domain-swapped dimer in the crystal structure, the oligomeric states of MtMce1A_36-148_ and MtMce4A_39-140_ were also determined by native mass spectrometry (MS) at two different concentrations (5 μM and 50 μM) to determine if there is a concentration-dependent dimerization. These studies further confirmed that both MtMce1A_36-148_ and MtMce4A_39-140_ are monomeric in solution at both the concentrations (*SI Appendix*, Fig. S6, S7).

**Table 2:** Molecular mass of MtMce1A and MtMce4A domains as calculated from SEC-MALS.

| Protein name | Theoretical monomeric molecular mass (kDa) | Protein-DDM conjugate (SEC-MALS) (kDa) | Protein (SEC-MALS) (kDa) | Empty DDM micelle (SEC-MALS) (kDa) |
| --- | --- | --- | --- | --- |
| MtMce1A<sub>36-148</sub> | 16.9 | - | 16.40 | - |
| MtMce1A<sub>38-325</sub> | 36.0 | 103.0 | 39.0 | 66.0 |
| MtMce1A<sub>126-454</sub> | 38.7 | 144.0 | 40.0 | 58.0 |
| MtMce1A<sub>38-454</sub> | 48.3 | 159.0 | 58.0 | 68.0 |
| MtMce4A<sub>39-140</sub> | 15.0 | - | 15.0 | - |
| MtMce4A<sub>39-320</sub> | 34.4 | 102.5 | 34.9 | 67.5 |
| MtMce4A<sub>121-400</sub> | 34.6 | 128.9 | 44.7 | 65.1 |
| MtMce4A<sub>36-400</sub> | 43.5 | 142.0 | 55.4 | 70.1 |

Interestingly, comparison of the secondary structure content of MtMce4A_39-140_ calculated from the CD spectrum with the crystal structure showed higher β sheet content (39.04%) in crystal than from the CD spectra (28.1%, *SI Appendix*, Table S4) indicating that the protein is more structured in the crystallization condition. Moreover, thermal melting analysis of MtMce1A_36-148_ and MtMce4A_39-140_ showed that they undergo heat-induced conformational changes (*SI Appendix*, Supplementary Results, Fig. S8 and S9). It is possible that in the purified conditions, both MtMce1A_36-148_ and MtMce4A_39-140_ are in non-native conformations and MtMce4A_39-140_ attains native conformation in the crystallization buffer. Therefore, the purified MtMce4A_39-140_ was exchanged in crystallization buffer and analyzed by SEC-MALS. Surprisingly, SEC-MALS analysis showed only the presence of monomeric MtMce4A_39-140_ also in the crystallization buffer (*SI Appendix*, Fig. S10). However, dimer formation was observed when MtMce4A_39-140_ was heated slowly, up to 50 °C in the crystallization buffer (0.7 M ammonium sulfate) (*SI Appendix*, Fig. S10). We speculate that the, incubation of this protein solution with the crystallization solution at 22 °C has probably facilitated the protein to attain a more compact and native state. The dimer could have selectively crystallized owing to better crystal contacts when compared to the monomer, indicating that the domain-swapped dimer is possibly a crystallization artifact.

### 2.5 *Comparison of* MtMce4A_39-140_ structure with *E. coli* and *A. baumannii* homologs

In spite of the initial non-native conformation of MtMce4A_39-140_, the overall MCE fold with a seven-stranded β-barrel is conserved in the MtMce4A_39-140_ crystal structure as also observed in the homologs from *E. coli* (EcMlaD, EcPqiB, and EcLetB) and *A. baumannii* MlaD (AbMlaD) (14–16, 18–20). As per our experimental data, the domain-swapped dimer formed by MtMce4A_39-140_ is most likely a crystallization artifact. Therefore, we used the compact folded monomer for structural analysis and comparison. The compact monomer is formed by residues 32 to 106 from chain A and residues 107 to 145 from chain B.

Superposition of the MCE domain from MtMce4A on EcMlaD and AbMlaD yields a root mean square deviation (RMSD) of 1.70 Å and 2.62 Å, respectively (Fig. 3*A*). The overall topology of the protein is conserved with conformational differences mainly in the loop regions and a few other secondary structural elements. For example, the β2a-52 to 54 is present only in the MtMce4A_39-140_ and not in EcMlaD and AbMlaD. The β4-β5 has an extra helix in MtMce4A_39-140_ and AbMlaD (45 residue insertion) and this helix is absent in the EcMlaD. The β5 strand and the hydrophobic β5-β6 loop (also referred as <u>P</u>ore <u>L</u>ining <u>L</u>oop; PLL) involved in forming the hydrophobic central pore, has a different conformation in MtMce4A_39-140_, which contrasts with the EcMlaD and AbMlaD. The β6-β7 loop in MtMce4A_39-140_ is a proline-rich loop whereas it is lined with charged residues in EcMlaD and AbMlaD. In addition, the density of the β6-β7 loop is missing in EcMlaD crystal structure and is present in MtMce4A_39-140_ and AbMlaD. The β7a-β7b is connected by a helix in MtMce4A_39-140_, by a loop in EcMlaD and β7a is absent in AbMlaD. The homologous β7b strand is much smaller in EcMlaD and AbMlaD compared to Mce4A_39-140_.

**Fig 3:**
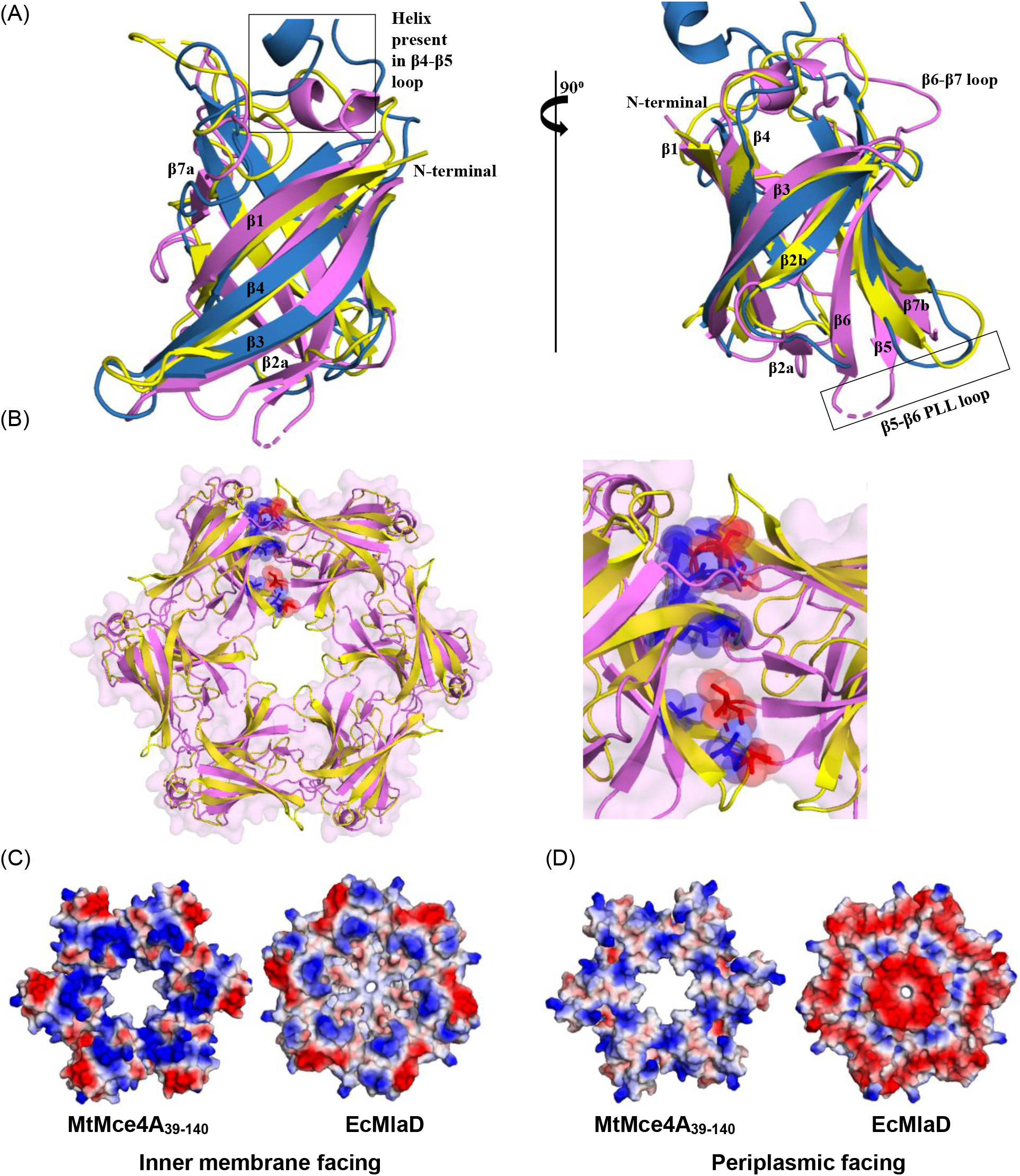
**(A)** Structural superposition of MCE domains of *Mtb* (MtMce4A_39-140_;pink), *E.coli* (EcMlaD; yellow) and *A.baumannii* (AbMlaD; blue). **(B)** Cartoon representation of the hypothetical homohexamer of MtMce4A_39-140_ (*pink*) generated based on EcMlaD homohexamer (*yellow*). The residues from two monomers (chains A and B) involved in the steric clashes are shown as sphere and sticks in blue and red colors. These clashes are between β2-β3 loop residues K61,Y62, R63 of chain A and β3-β4loop residues S76, G77, Q79 of chain B; between β5 strand residue A103 of chain A with β5-β6 loop residue I107 of chain B; between β6 strand residue E114 of chain A with β5-β6 loop residue A50 of chain B and between β7 strand residue L140 of chain A with β5-β6 loop residue T106 and I107 of chain B. **(C)** Electrostatic potential surface (inner membrane facing side) of the hypothetical homohexamer of MtMce4A_39-140_ (left) and EcMlaD homohexamer (right). The inner membrane facing sides are comparable with each other with high positive charges. **(D)** Electrostatic potential surfaces (periplasmic facing side) of the hypothetical homohexamer of MtMce4A_39-140_ (left) and EcMlaD homohexamer (right). The periplasmic side of MtMce4A_39-140_ has more positively charged residues, which is in contrast to the highly negative surface of EcMlaD.

Similarly, MtMce4A_39-140_ was superposed with MCE domains of EcPqiB1-3 and EcLetB1-7 monomers (*SI Appendix*, Fig. S11). The superposition showed that the β-barrel fold is conserved, and the observed differences are mainly in the loop regions. For example, the β2a-52 to 54 is unique in MtMce4A_39-140_ and is absent throughout in EcPqiB1-3 and EcLetB1-7. The β3-β4 loop conformation present on the exterior surface varies amongst MtMce4A_39-140_, EcPqiB1-3 and EcLetB1-7. Noteworthy, the length of β3-β4 loop remains constant (4 residues) in all the MCE domains except the EcPqiB3, which has 18 residues in the loop. The PLL (β5-β6 loop) comprising the hydrophobic channel is much longer (16-27 residues) in EcPqiB1-2 and EcLetB1-7 when compared to the MtMce4A_39-140_ (5 residues). We found that the PLL in EcPqiB3 has only 7 residues and it is the only MCE domain amongst EcPqiB and EcLetB, which share this feature with MtMce4A_39-140_. Interestingly, the conformation of all the PLL varies throughout the MCE domains of EcPqiB1-3 and EcLetB1-7 (*SI Appendix*, Fig. S11). Furthermore, the β6-β7 loop in MtMce4A_39-140_ has a different conformation as compared to EcPqiB1-3 and EcLetB1-7. Amongst the available Mce SBP structures and MtMce1/4A-F, MtMce4A_39-140_ has maximum number of proline residues in the β6-β7 loop. The role of this proline rich loop is still not understood.

The monomeric nature of MtMce4A_39-140_ is contrasting to the other Mce proteins (MlaD, PqiB, LetB) from *E. coli* and *A. baumannii*, which forms a homohexamer (14–16, 18–20). Based on the EcMlaD, we generated a hypothetical homohexamer of MtMce4A_39-140_ (Fig. 3*B*). Interestingly, the domain interface of the hypothetical homohexamer of MtMce4A_39-140_ has multiple steric clashes, which will preclude the formation of homohexamers in MtMce4A (clashes between chain A and B are shown in Fig. 3*B*). These clashes are absent in EcMlaD, AbMlaD, EcPqiB1-3, and EcLetB1-7, where homohexamers have been formed.

### 2.6 Elongated conformation of MtMce1A_36-148_ and MtMce4A_39-140_ in solution

SAXS experiments were performed to gain information on the shape and size of the purified MtMce1A_36-148_ and MtMce4A_39-140_. Measured intensities, *I*(*q*), are displayed as a function of the modulus, *q*, of the scattering vector. Structural parameters calculated from the scattering intensities are given in the Table S5. The radius of gyration (*R*_*g*_) and maximum interatomic distances (*D*_*max*_) were determined to be 21.6 Å and 70 Å for MtMce1A_36-148_, and 21.7 Å and 80 Å for MtMce4A_39-140_, respectively (*SI Appendix*, S12). Interestingly, the determined *D*_*max*_ of both MtMce1A_36-148_ and MtMce4A_39-140_ is much higher than the maximum diameter of homologous EcMlaD (35 Å) pointing towards an elongated structure for both the proteins. Further, *ab initio* molecular shape reconstructed by DAMMIN indicate that both MtMce1A_36-148_ and MtMce4A_39-140_ attains an elongated shape in the purified conditions (Fig. 4*A* and *B*). From SEC-MALS and SAXS, we know that the proteins exist as monomers in solution. Therefore, the *ab initio* shape of MtMce4A_39-140_ was fitted with the compact and elongated monomers of MtMce4A_39-140_. The agreement in terms of reduced chi-squared (χ^2^) of compact and elongated model calculated against the experimental SAXS data was 10.0 and 2.0, respectively (Fig 4B). Similarly, in case of MtMce1A_36-148_, a template-based model (obtained from Robetta) was used for fitting the SAXS data and the χ^2^ for compact and elongated model were 14.0 and 11.0, respectively (Fig 4A), also here favoring slightly the elongated model. Further, the domain-swapped region of MtMce1A_36-148_ was optimized by rigid-body refinement and this improved the χ^2^ to 4.2. In summary, the elongated models fit relatively better than the compact model in both cases, suggesting that both MtMce1A_36-148_ and MtMce4A_39-140_ attains an elongated shape in the purified conditions.

**Fig. 4:**
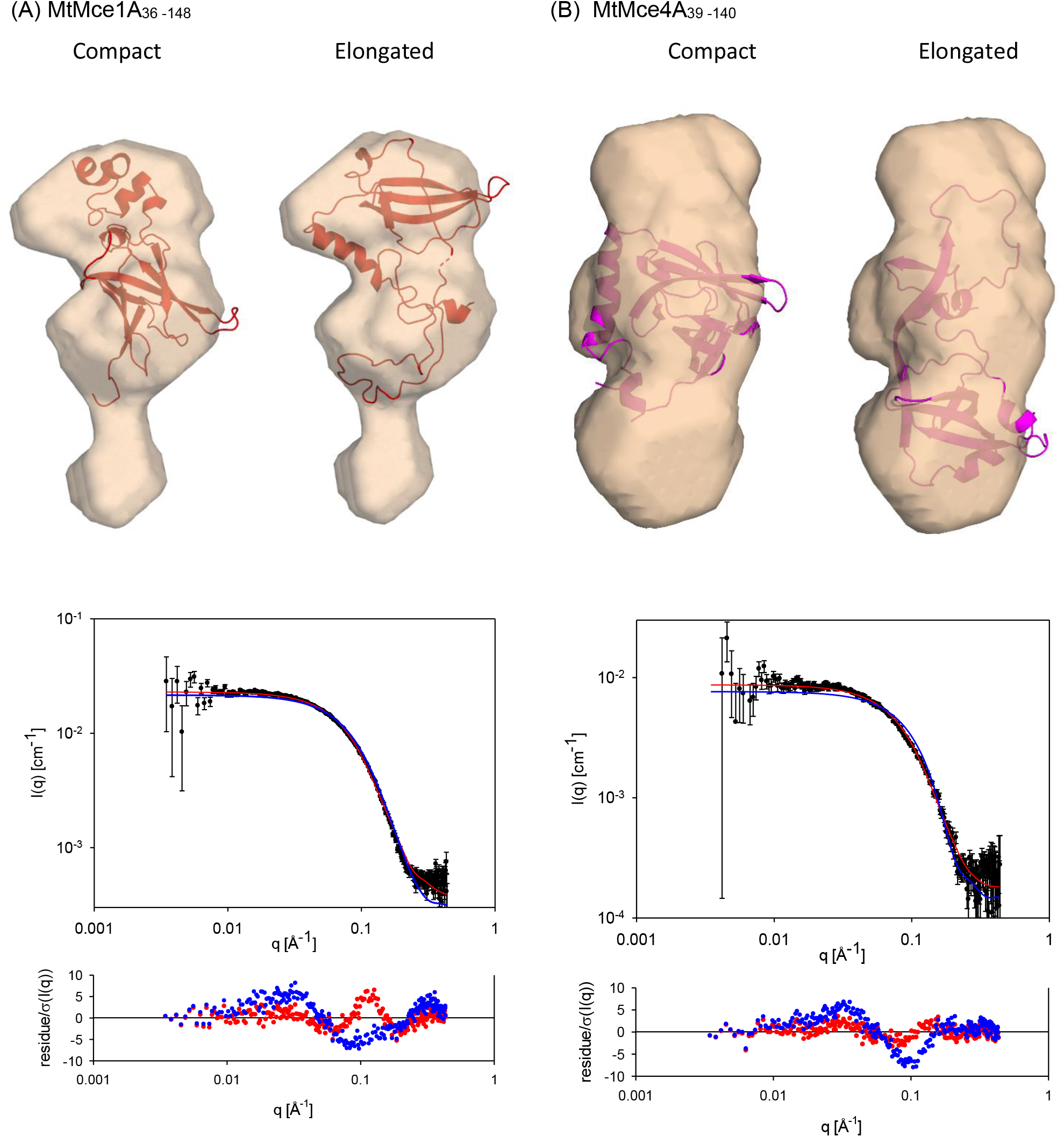
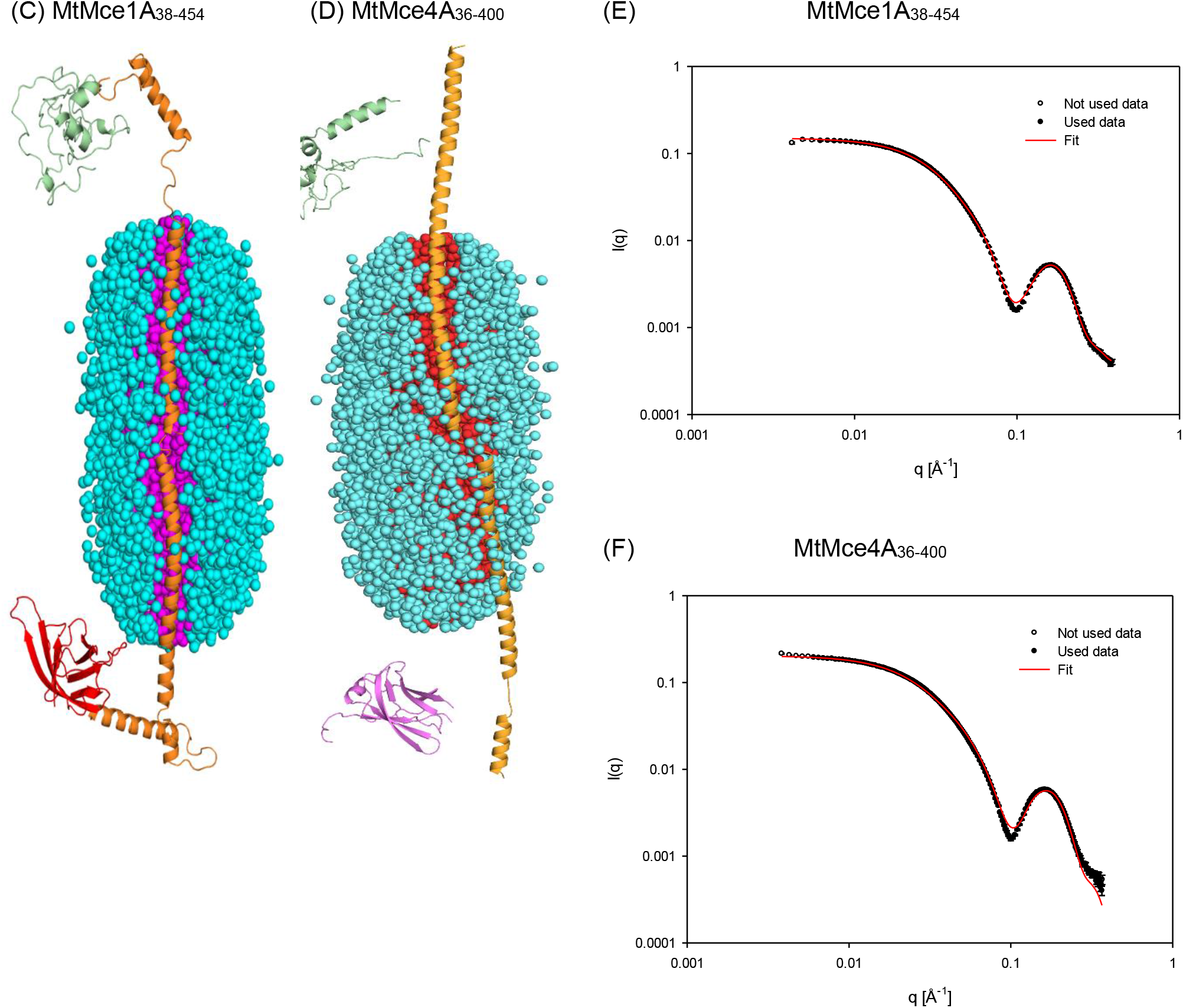
**(A)** The *ab initio* shape generated by DAMMIN for MtMce1A_36-148_ superposed on the elongated (left) and compact (right) monomeric models of MtMce1A_36-148_. The corresponding fits of the experimental SAXS data (black) with the elongated (red) and compact (blue) monomers are shown below. **(B)** The *ab initio* shape generated by DAMMIN for MtMce4A_39-140_ superposed on the elongated (left) and compact (right) monomeric models of MtMce4A_39-140_. The corresponding fits of the experimental SAXS data (black) with the elongated (red) and compact (blue) monomers are shown below. **(C)** The model of MtMce1A_38-454_ and **(D)** MtMce4A_36-400_ interacting with the core of DDM micelle.The DDM molecules are represented as spheres and the protein as cartoon. **(E)** The fit (red line) of experimental SAXS data (*black dots*) with the proposed models of MtMce1A_38-454_ and **(F)** MtMce4A_36-400_.

### 2.7 Helical domain of MtMce1A and MtMce4A interacts with DDM core

In addition to the soluble MtMce1A_36-148_ and MtMce4A_39-140_, SAXS measurements were also performed for the other MtMce1A and MtMce4A domain constructs in SEC-inline mode as they have been purified in the presence of DDM (*SI Appendix*, Table S6 and S7). The elution profile has two peaks, one for the PDCs and one for the empty micelles. The SAXS scattering data of the PDCs display a minima at a scattering vector modulus of 0.1 Å^−1^ followed by a broad bump, which reconfirms the presence of nearly intact detergent micelle together with the protein. To further analyze the SAXS data, models of MtMce1A and MtMce4A were generated and optimized as described in the methods section (*SI Appendix*). In both MtMce1A and MtMce4A models, the helix turns back at residues E248 and D215, respectively, to form a coiled-coil structure, where the coiled-coil helices are held together by hydrophobic interactions and which brings the tail domain closer to the MCE domain. These models are referred as ‘‘coiled-coil models’’. A second variation of this model was also generated by opening the helical domain to form an extended helix keeping the tail domain far away from MCE domain. This model is referred as “extended helical model” (Fig. 4*C* and *D*).

With our experimental data, it is clear that the MCE domain is soluble and the presence of helical domain required detergent for the purification. Therefore, the detergent micelle has to interact with the helical domain. Although the helical domain has a high number of hydrophobic amino acids, it also has polar residues which precludes the possibility of the helix completely inserted into the core of the micelle like a typical transmembrane protein. Furthermore, calculations of the SAXS intensity with the helix inserted into the core, show that SAXS intensity for these models smears out the minimum, so that it is not as deep as in the data. Therefore, a core-shell model of the detergent micelle was used where the helical domain is interacting with the surface of the core (dodecyl chains) of the micelle (21). On testing multiple micelle shapes, the best fit was obtained when using a superellipsoid shape with the long axis along the helical domain which maximizes the interaction of the helix with the core of the micelle. The micelle size (aggregation number) was initially estimated from SEC-MALS and SAXS scattering analysis (21) to be in the range of 125-200. However, in cases where the fits were not satisfactory, it was further varied in a reasonable range for obtaining the best fit to the SAXS data.

With these assumptions, both the coiled-coil as well as extended helical models for each of the MtMce1A and MtMce4A constructs were optimized (10 independent runs) together with the micelle with appropriate aggregation number to fit the SAXS data. For MtMce1A_38-325_, as well as MtMce4A_39-320_ (MCE+Helical domains), both, the coiled-coil and extended, models showed convincing fits with χ^**2**^ values ranging from 3-20. In case of MtMce1A_38-325_, the best fit to the data are very good with the low *q*, minimum and broad bump being very well reproduced (*SI Appendix*, Fig. S13). For MtMce4A_39-320_, the best fit is very good in the whole range for the extended model, however, with a tendency that the model structure is slightly too large and the minimum in the model curve being slightly higher than the data (*SI Appendix*, Fig. S14). In contrast, for the best fit with the coiled-coil structure, the size of the model seems slightly too small and there are more systematic deviations around the minimum and bump (*SI Appendix*, Fig. S14). From the χ^**2**^ values and these observations, it can be concluded that the extended model is best in terms of reproducing the data.

In case of MtMce1A_38-454_ (MCE+Helical+Tail domains), the extended helical model (Fig. 4C) has a better fit with a χ^**2**^range of 15-24 compared to the coiled-coil model (χ^**2**^=22-50). The extended model fits the data well in the full *q* range with a small deviation around the minimum, where the model curve is not quite low enough (Fig. 4E). The coiled-coil model is too small with some deviations at low *q*, and the optimization compensates partly by displacing the MCE domain away from the Helical+Tail domain, leading to some disconnectivity of the structure (*SI Appendix*, Fig. S15). In case of MtMce4A_36-400_, the data are not fitted well at the high *q* values for both models, although, the low *q* data fits better in the extended model, favoring the extended helical model (Fig. 4D). Similar to the MtMce1A_38-454_ coiled-coil model, the MCE domain and the Helical+Tail domain gets disconnected in the MtMce4A_36-400_ coiled-coil model favoring the extended model (Fig. 4F and *SI Appendix*, S15).

The Helical+Tail domain fits for MtMce4A_121-400_ extended and coiled-coil models have similar χ^**2**^ values and are in the range of 4.6-8.0. The extended model fits very well in the high and low *q* range and the coiled-coil has deviations at the high *q* data (*SI Appendix*, Fig. S16). Both these models have less deep minima with respect to the data. The MtMce1A126-454 extended model fitting has a χ^2^ range of 40-75, whereas, the coiled-coil model fit shows a χ^2^ between 114-182. Similar, to the MtMce4A_121-400_ models, the minima is also less deeper in MtMce1A_126-454_ models. It could also be concluded that the extended models fit better in case of Helical+Tail domain (*SI Appendix*, Fig. S17). Additionally, we have to accept that the tail domain is unstructured with greater uncertainty in the structure prediction. This could also be a reason for poorer SAXS fits for all the constructs with the tail domain. The counting statistics of the data for the samples varies somewhat and therefore χ^**2**^values also varies, it is observed that the χ^**2**^ values are often higher for data with good counting statistics. Therefore, we decided also to calculate R factors and weighted R factors as used in crystallography. R factors are dominated by the high intensities at low q, whereas weighted R factors are a normalized measure of (χ^**2**^)^0.5^. The determined values are both in the range 1-5 % (*SI Appendix*, Table S6, S7). They reveal that the deviation between data and fits are more similar than the χ^**2**^ values suggest.

Considering the low resolution information in the SAXS data as well as the possible errors in the generated MtMce1A and MtMce4A models, which are partly based on structural predictions, our analysis gives the best possible explanation for the observed SAXS data in a qualitative and in a semi-quantitative manner. Among the coiled-coil and extended helical models, the fit for the extended helical models are relatively better overall. It is possible that the extended helical model is more favorable because the DDM micelle can interact with a large area of the helix in the extended helical model as compared to the smaller area in the coiled-coil model.

## 3 Discussion

We have classified the *Mtb* Mce1A-1F and Mce4A-4F SBPs into four different domains based on the secondary structure prediction, which shows the presence of a unique tail domain in SBPs from *Mtb*. Further characterization shows that the full length as well as all the domains of MtMce1A and MtMce4A remain as monomers in solution when purified individually. Moreover, only the MCE domain is soluble whereas the MCE+Helical and the Helical+Tail domains require detergents for their solubility. Further, SAXS analysis of MtMce1A, MtMce4A and their domains suggests that the individual protein may adopt the ‘extended helical’ model where the MCE and tail domains are far away from each other. Also, structural analysis of MtMce4A_39-140_ suggests that the homohexamer cannot be formed at least in Mce4A due to multiple steric clashes. It is possible that in case of *Mtb*, the six MceA-F SBPs can interact with each other to form heterohexamers rather than homohexamers. Taken together, on the basis of all these data, we propose a model for the *Mtb* Mce-complex, where the MceA-F SBPs would exist as an “extended helical heterohexamer” (Fig. 5). In this model, the MCE domain from each SBP will form a ring like structure and the long helical domain will form a hollow channel. Inferring from the SAXS data, we speculate that the space occupied by the shell of the DDM micelle on the helical domain of MtMce1A and MtMce4A would be replaced by the interacting MceB-F SBPs.

**Fig. 5:**
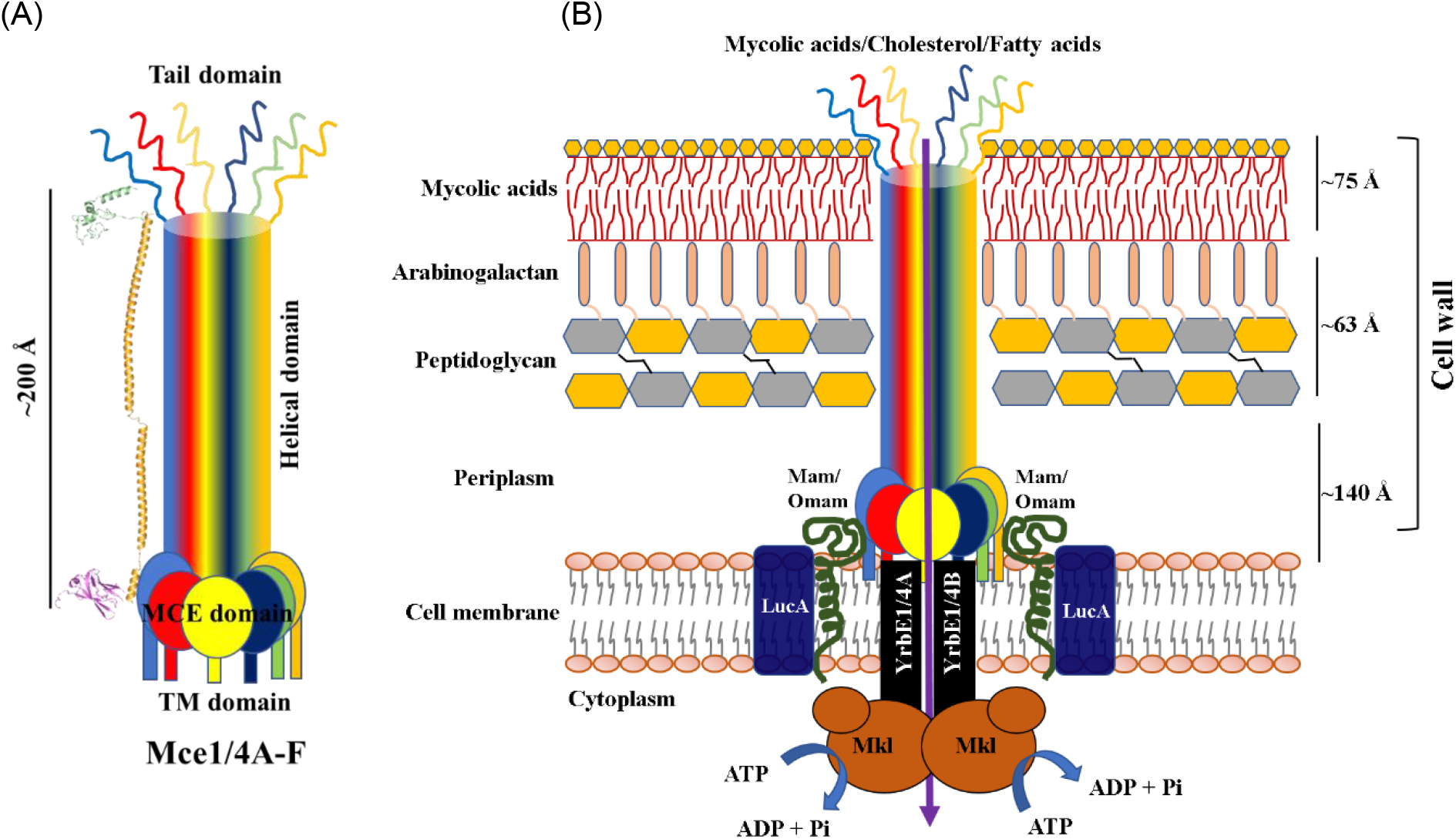
**(A)** Proposed heterohexameric arrangement of *Mtb* Mce1/4A-F SBPs. The model for MtMce4A is represented in cartoon. **(B)** Schematic representation of Mce1/4 complex inside the cell wall of *Mtb*. The Mce1/4A-F SBPs would be embedded in the outer lipid layer, where the soluble N-terminus would be predicted to remain in the periplasmic space. The proposed permeases YrbE1/4AB would encompass the inner cell membrane connecting the Mce1/4A-F with Mkl. The Mkl would act as an ATPase and would convert the energy of ATP for the transport of lipid molecules across the membranes. The Mam/Omam and LucA would act as accessory proteins, involved in stabilizing the entire complex.

Additionally, the core of the DDM micelle could correspond to the hollow hydrophobic channel formed by the six interacting MceA-F SBPs in the physiological condition. The length of *Mtb* periplasmic space is ~140 Å (22) and the calculated *D*_*max*_ from SAXS for the MCE+Helical+Tail domains is approximately 200 Å. Therefore, we propose that the MCE domain would be present in the periplasm, the helical domain would extend from the periplasm to the mycolic acid layer and the tail domain would extend to the surface of the cell wall. The tail domain reaching the surface of the cell wall could play a role in the substrate recognition, binding and transport of lipids into the ‘channel’. Although the individual tail domains of MceA-F SBPs are predicted to be unstructured, it is possible that they acquire some structure upon interaction with each other. In the inner membrane side, the MCE domain heterohexamer would be connected to the YrbEAB permeases, which will form the transmembrane channel. The Mkl (MceG) interacting with the YrbEAB in the cytoplasmic side of the inner membrane would act as the driver of the transport in an ATP-dependent manner. Additionally, the Mam proteins and LucA would act as stabilizing proteins (23, 24) to form the Mce complex. Therefore, the Mce complex would form a direct connection between the cytoplasm and cell surface facilitating the transport of lipids across the cell wall (Fig. 5).

The central pore of all the reported Mce SBP hexamers are comprised of highly hydrophobic residues also known as PLL, which allows the transport of small hydrophobic lipid molecules across the membranes. The variation in the length of PLL depends on the transport mechanism followed by the particular Mce complex. For example, EcMlaD and AbMlaD has a smaller PLL (6 residues) because they follow a ferry-based transport mechanism. Whereas, the PLL is longer in EcPqiB and EcLetB (17-27 residues) in agreement with that they follow a tunnel-based lipid transport mechanism. It has been reported that the PLL of EcMlaD, AbMlaD, EcPqiB_1-3_, and EcLeTB_1-7_ follows a pattern of ‘‘ϕxxϕϕ’’, where ‘ϕ’ denotes hydrophobic amino acid and ‘x’ represent any amino acid (16). This pattern has been followed in MtMce1A (_112_ATTVF_116_) and MtMce4A (_104_GNTIF_108_), but it does not align with other Mce SBPs from *Mtb* (*SI Appendix*, Fig. S18). Instead the other *Mtb* Mce SBPs follow a pattern of ‘‘xxxϕϕ’’in the PLL. This conserved “duo” of consecutive hydrophobic residues in the central pore of MtMce4A-4F SBPs is indicating towards formation of a heterohexameric channel. In addition, the helical domain of the MtbMceA-F SBPs also has high number of hydrophobic residues although a clear ‘motif’ is not observed.

The inner membrane-facing surface of homohexamers of EcMlaD, AbMlaD and 1^st^ ring of EcPqiB and EcLetB has predominantly positively charged residues, which would interact with the negatively charged outer leaflet of the plasma membrane. The hypothetical MtMce4A_39-140_ homohexamer also share the same surface electrostatics on the inner membrane facing side (Fig. 4*C*). The periplasmic side of the EcMlaD, AbMlaD homohexamer is lined with negatively charged residues, which are needed for the hydrophilic interactions. In contrast, the periplasmic side of MtMce4A_39-140_ is positively charged and is more comparable to the 3^rd^ ring of EcPqiB (Fig. 4*C*). EcPqiB_3_ and MtMce4A_39-140_, both are connected to a long helix and we predict that this helix is shielding the interaction of MCE domain with the periplasm. With the current knowledge of *Mtb* Mce SBPs and its comparison with homologous Mce SBPs, we propose that the lipid transport mechanism in *Mtb* Mce-complex would be tunnel-based as observed in EcPqiB and EcLetB. We believe that the main difference may be the involvement of the tail domain in substrate recognition and shuttling of the lipids from the outer surface of the *Mtb* whereas, the shuttling in EcPqiB and EcLetB probably involves other outer membrane proteins. Indeed, a higher resolution structure of MtMceA-F SBPs or the entire Mce complex is needed to further understand the detailed structural arrangement as well as the lipid transport mechanism of the mycobacterial Mce complexes.

## 4. Materials and Methods

Details of plasmids, constructs, protein expression, purification, experiments including SEC-MALS, SAXS, crystallization, CD, native MS, along with sequence and structural analysis are provided in *SI Appendix*.

## Author Contributions

P.A. and D.S. performed the work on Mce4A-4F and Mce1A-1F proteins, respectively. P.A. and R.V. determined the crystal structure of Mce4A domain. J.S.P. performed the SAXS analysis and simulations for Mce1A and Mce4A domains. M.J.H., R.S. and A.V.M. assisted in the cloning and/or purification of Mce1C & Mce1E, Mce1B and Mce1D, respectively. M.L. and J.J. conducted the native mass spectrometry experiments. L.W.R contributed in the discussion, manuscript writing and co-supervision. P.A., D.S., and R.V. designed the experiments, analyzed the data and wrote the manuscript with comments from all authors. R.V. conceptualized and supervised the work and acquired funding.

## Acknowledgments

This work was supported by funding from Jane and Aatos Erkko foundation, Sigrid Juselius Foundation, I4Future doctoral program (MSCA-COFUND by Horizon 2020 European Union, grant agreement number 713606), Academy of Finland (332967) and Biocenter Oulu. R.V. also acknowledges Academy of Finland for the Academy Research Fellowship (319194). MaxIV, DLS, ESRF and iNEXT (funded by Horizon 2020 European Union, grant agreement number 653706 and 730872) are acknowledged for their support in X-ray and SAXS data collection. We acknowledge the support of Biocenter Oulu (proteomics and protein analysis, structure biology and DNA sequencing core facilities) and Biocenter Kuopio/FINStruct (native mass spectrometry). The FT-ICR MS facility is also supported by the European Regional Development Fund (A70135). We acknowledge the support of Dr. Hongmin Tu for CD analysis.

## Database Depositions

The X-ray structure for SeMet-MtMce4A_39-140_ and MtMce4A_39-140_ has been deposited in the PDB with the accession codes 7AI2 and 7AI3, respectively. The SAXS structures for MtMce1A and MtMce4A domains are deposited in SASDB. The accession codes for MtMce1A_36-148_, MtMce1A_38-454_, MtMce1A_126-454_, and MtMce1A_38-325_ are SASDJU9, SASDK32, SASDK22, and SASDJZ9, respectively. The accession codes for MtMce4A_39-140_, MtMce4A_36-400_, MtMce4A_121-400_, and MtMce4A_39-320_ are SASDJV9, SASDJW9, SASDJX9, and SASDJY9.

## Notes

### Competing Interest Statement

The authors have declared no competing interest.

### Summary of Updates

SASDB codes updated

