## SupplementaryInformation for "Structural insights on the substrate-binding proteins of the *Mycobacterium tuberculosis* mammalian-cell-entry (Mce) 1 and 4 complexes"

**Key words**

**Abbreviations**

Ab- *Acetinobacter baumannii*

CD- Circular dichroism

$C_{12}E_9$- Dodecyl nonaethylene glycol ether

DDM- n-Dodecyl β-D-maltoside

Ec- *Escherichia coli*

FC-12- Fos choline-12

Hepes- 4-(2-Hydroxyethyl)piperazine-1-ethanesulfonic acid

IM- Inner membrane

IPTG- Isopropyl β-D-thiogalactopyranoside

MCE- Mammalian-cell-entry

MES- 2-(N-morpholino) ethanesulfonic acid

MlaD- Membrane lipid asymmetry

PDC- Protein detergent complex

PqiB- Paraquat-inducible protein B

LetB- Lipophilic envelope spanning tunnel B

MS- Mass spectrometry

Mtb- *Mycobacterium tuberculosis*

OM- Outer membrane

SAD- Single-wavelength anomalous dispersion

SBP- Substrate-binding proteins

SAXS- Small angle X-ray scattering

SEC- Size exclusion chromatography

SeMet- Selenomethionine

Tb- Tuberculosis

Tris- Tris(hydroxymethyl)aminomethane

### Materials and Methods

#### *Biochemicals*

The genomic DNA of *Mtb H37Rv* was purchased from ATCC. *Phusion* DNA polymerase and restriction enzymes used for cloning were purchased from Thermo Scientific (Massachusetts, USA) and New England BioLabs. The Ni-NTA chromatography resin was obtained from Qiagen (Hilden, Germany).

#### *Cloning, expression and purification of MtMce1A-1F and MtMce4A-4F*

Individual MtMce1A-1F and MtMce4A-4F genes were PCR amplified using *Mtb H37Rv* genomic DNA as the template with specific primers (Table S8, S9). Each amplicon was cloned into pETM11 vector (EMBL) using restriction based cloning method resulting in an N-terminal $His_6$-Tag followed by a TEV protease site, *MceA-F* gene and C-terminal $His_6$ Tag. For protein expression, the plasmid was transformed into *E. coli* BL21-RIPL competent cells. Overnight cultures were grown at 30 ºC until $O.D_{600}$ reached 0.6, and the expression of the protein was induced with 0.5 mM IPTG (isopropyl β-D-thiogalactopyranoside) at 16 ºC overnight. Cells were harvested by centrifugation at 4000 g. The bacterial pellet was resupended in the desired lysis buffer with a suitable detergent (Table S10). Cells were lysed using sonicator (30% amplitude, 3 min) and the lysate was centrifuged at 15000 g at 4 ºC for 30 min. The supernatant was then filtered (0.45 µm; Millipore), and proteins were allowed to bind to $Ni^{2+}$-NTA matrix for 1 hour. The beads were washed, and bound proteins were eluted from the Ni-NTA column using 400 mM of Imidazole in the desired elution buffer (Table S10). At this step the concentration of the detergent was reduced to 5 mM. The eluted protein was analyzed on a 12% or 18% SDS-PAGE and was concentrated (spin concentrator, molecular cut-off: 30 kDa, Millipore) and injected on a SEC column (Superdex 200 10/300 or Superdex 75 HiLoad 16/600 column, GE-Healthcare). The identity of the eluted protein was further confirmed with peptide mass finger printing analysis.

#### *Expression and purification of MtMce1A and MtMce4A domains*

Based on the secondary structure analysis, MtMce1A and MtMce4A domain constructs were generated. They were cloned in pETM11 using restriction free cloning methods: the constructs were named according to the secondary structural features: 1) MCE domain ($MtMce1A_{36-148}$ and $MtMce4A_{39-140}$), 2) MCE+Helical+Tail domain ($MtMce1A_{38-454}$ and $MtMce4A_{36-400}$), 3) Helical+Tail domain ($MtMce1A_{126-454}$ and $MtMce4A_{121-400}$) 4) MCE+Helical domain ($MtMce1A_{38-325}$ and $MtMce4A_{39-320}$) and 5) Tail domain ($MtMce4A_{321-400}$). The expression and purification protocols were similar to the corresponding full-length proteins. Only the MCE domain constructs ($MtMce1A_{36-148}$ and $MtMce4A_{39-140}$), were soluble without the use of detergents and different buffer was used for lysis and elution (Table S10).

For Selenomethionine (SeMet) labelled $MtMce4A_{39-140}$, the construct was transformed to auxotrophic strain of *E. coli* (B834) and grown according to the protocol from Molecular Dimension (1). The expression and purification protocols were similar to the native $MtMce4A_{39-140}$. SeMet incorporation was confirmed by electrospray ionization liquid chromatography-mass spectrometry (ESI LC-MS) which showed 100% incorporation of SeMet into the protein.

***SEC-MALS of Mce1A and Mce4A domains***

SEC-MALS analysis of purified MtMce1A and MtMce4A domains was carried out using SEC coupled the miniDAWN$^{TM}$ TREOS (*Wyatt Technologies)*. Approximately 5-6 mg ml$^{-1}$ of purified protein was loaded on the pre-equilibrated Superdex 200 10/300 using an autosampler and was run at a rate of 0.4 ml min$^{-1}$ at 4 °C in a Shimadzu HPLC/FPLC system. The samples were then passed through a refractive index (RI) detector and subsequently through the MALS detector. The cumulative data collected for UV, MALS and RI, were analyzed using the *ASTRA* software (*Wyatt technologies*). The proteins were bound to detergent; therefore, protein-conjugate analysis method was incorporated for analysis. The detergent was considered as a modifier and the recommended dn/dc values of 0.1473 ml g$^{-1}$ for n-Dodecyl β-D-maltoside (DDM) were used for the protein-conjugate analysis. The soluble MtMce1A$_{36-148}$ and MtMce4A$_{39-140}$, analysis was done without using the protein conjugate template.

For understanding the effect of heat and higher ionic strength on the oligomeric state of the protein, purified MtMce4A$_{39-140}$ was subjected to buffer exchange (0.1 M MES [2-(N-morpholino)ethanesulfonic acid], 0.7 M ammonium sulfate, pH 6.0) using a 10 kDa molecular mass cut-off amicon concentrator. Mce4A$_{39-140}$ was heated to 50 °C in a thermocycler, with initial one min incubation at 20 °C followed by 0.8 °C increase per min up to 50 °C and a final incubation at 50 °C for one min. The heated protein was then centrifuged at 10000 g for 5 mins and the supernatant was injected to Superdex 200 10/300 column pre-equilibrated with a buffer containing 0.1 M MES and 0.7 M ammonium sulfate at pH 6.0 coupled to a MALS detector. The results were processed using both UV and RI signals as the source of concentration to calculate the molecular mass.

***Circular Dichroism (CD) spectroscopy of MtMce1A and MtMce4A* domains**

The MtMce1A and MtMce4A domains were diluted in water to obtain a lower buffer and salt concentration. The protein concentration used for CD measurements (Chirascan CD spectrophotometer, Applied Photophysics, Surrey, U.K.) was 0.05 mg ml$^{-1}$. The secondary structure analysis of MtMce1A and MtMce4A domains purified with DDM and without DDM using CDNN and BestSel softwares, respectively (18, 19). For the determination of the thermal melting temperature ($T_m$), the sample was heated at a rate of 1 °C per min from 22 °C to 92 °C. The melting curves were calculated by comparing the spectra from 190 to 280 nm with the global fit analysis protocol as implemented in the Global3 software.

**Native mass spectrometry of MtMce1A$_{36-148}$ and MtMce4A$_{39-140}$**

MtMce1A$_{36-148}$ and MtMce4A$_{39-140}$ were buffer-exchanged to 20 mM ammonium acetate (pH 6.8) using PD Miditrap G-25 columns (GE Healthcare, Sweden). Mass spectra were measured on a 12-T Bruker solariX XR FT-ICR mass spectrometer, using Apollo-II electrospray ion source (Bruker Daltonics, Bremen, Germany). The instrument was calibrated using sodium perfluoroheptanoic acid (NaPFHA) clusters and operated with ftmsControl 2.2 software. The mass spectra were further analyzed using DataAnalysis 5.1 software.

***SAXS of MtMce1A and MtMce4A domains***

*Data collection:* SAXS data for the purified Mce1A and Mce4A domains were collected on the B21 beamline of Diamond Light Source (DLS), UK. Data was collected based on the standard protocols for inline-SEC SAXS and batch mode measurement using a Pilatus 2M two-dimensional detector at a sample-detector distance of 4.014 m and at a wavelength of λ = 0.99 Å. Inline SEC-SAXS measurements were collected for domains purified in the presence of detergent DDM (MtMce1A$_{38-325}$, MtMce1A$_{126-454}$, MtMce1A$_{38-454}$, MtMce4A$_{39-320}$, MtMce4A$_{121-400}$, and MtMce4A$_{36-400}$) at an initial concentration of 5 mg ml$^{-1}$. Batch mode measurements were collected for MtMce1A$_{38-148}$, and MtMce4A$_{39-140}$ at 2 mg ml$^{-1}$ and 1 mg ml$^{-1}$ respectively, with bovine serum albumin (BSA) as a control. For each batch mode concentration, 25 frames were collected.

*Data processing:* Data processing and analysis was done using the ScÅtter and ATSAS software packages (2). The 2D data were averaged to give a 1D data set of intensity, $I(q)$, vs $q$, where $q$ is the modulus of the scattering vector. The scattering of the buffer was subtracted from the protein scattering using the ScÅtter program. The data were rebinned using home-written software to be approximately equidistantly spaced on a logarithmic $q$ scale. The radius of gyration ($R_g$), forward scattering $I$ (0) and maximum particle distance ($D_{max}$) were calculated using PRIMUS. The hydrated volume $V_p$ was computed from the Porod invariant. The molecular weight was calculated based on two methods: Volume of correlation (3) and SAXSMoW (4). *Ab initio* shape was generated using DAMMIN (5). For MtMce4A$_{39-140}$, the compact monomer was generated by residues 32 to 106 from chain A and residues 107 to 145 from chain B of the crystal structure. The elongated monomer corresponds to the Chain B of the crystal structure. These were further provided as a template in Robetta to add the missing residues (6, 7). Whereas, for MtMce1A$_{36-148}$, the entire compact and elongated models were generated with Robetta using the MtMce4A$_{39-140}$ compact and elongated crystal structures as the template. The models were evaluated against the experimental data using a home-written program (8, 9). The helical and tail domains for MtMce1A$_{38-325}$, MtMce1A$_{126-454}$, MtMce1A$_{38-454}$, MtMce4A$_{39-320}$, MtMce4A$_{121-400}$, and MtMce4A$_{36-400}$ were generated using *iTasser*. A summary of the data collection and analysis parameters are summarized in supplementary Table 3 and 4.

*Detergent and protein model fitting for* MtMce1A$_{38-325}$, MtMce1A$_{126-454}$, MtMce1A$_{38-454}$, MtMce4A$_{39-320}$, MtMce4A$_{121-400}$, and MtMce4A$_{36-400}$*:* The SAXS data for the complexes of DDM and the various constructs were also analyzed using home-written software. The program is based on the methods described in (8-11). The DDM micellar structure is represented by Monte Carlo points in a tri-axial core-shell structure with super-ellipsoidal shape with shape parameter $t$ = 3 (12), and the protein is represented by the atoms in the PDB structures. When the protein overlaps with the core-shell structure, the corresponding Monte Carlo points were removed. The volume of the core was estimated from the number of points and the point density, and the aggregation number was calculated by dividing the core volume by the volume of a C12 chain (353 Å$^3$). The shell contains both DDM headgroups and solvating buffer, and the thickness of the shell was fixed at 10 Å. In practice, the aggregation number was kept fixed and the length of the long axis and of one of the short axes optimized, whereas the length of the third axis was calculated from these two and the aggregation number. The Monte Carlo points were assigned an excess scattering length corresponding to the electron densities of C12 tails and heads for points in the core and in the shell, respectively, taking into account the glycerol content of the buffer. Similarly, the excess scattering length of the atoms of the protein was adjusted taking into account the glycerol. The scattering of a hydration layer was added to the protein in the places where it is not in contact with the micelle. The protein structure was divided into three domains, namely MCE, Helical and Tail domains, to allow rigid-body refinement. The domains (MCE+Helical, Helical+Tail, and MCE+Helical+Tail, respectively for the three constructs) were connected by soft restraints as described in Vilstrup et al (9). The algorithm for generating the micelle including the estimates of the excess scattering length were checked by fitting a data frame from pure micelle from the elution profile, and it gave a satisfactory fit.

The SAXS data for all constructs have a deep minimum around $q$ = 0.1 $Å^{-1}$, followed by a pronounced secondary maximum. This behavior is qualitatively very similar to that of pure DDM micelles, and the first tests revealed that such a $q$ dependence could not be obtained when the protein penetrates significantly into the core of the micelles. Further tests showed that reasonable agreement with the SAXS data was obtained when the helix of the protein was along the long axis of the DDM micelle. Therefore, starting structures with this position were used in the optimizations. Additionally, a soft restraint that keeps the helix in contact with the micelle was introduced. The structure was optimized by random searches, initially with large amplitudes, which were gradually decreased during the optimization (9) or each structure, 10 independent runs were performed, each with 4000 cycles of optimization. The structure with the best agreement with the SAXS data in terms of reduced chi-squared, $\chi^2$, was selected as the resulting structure. Initially the aggregation numbers were estimated from the SEC-MALS results, however, in some cases, this did not give good fits to the SAXS data. Therefore, the aggregation number was varied in a reasonable range for these cases.

***Crystallization, data collection and structure determination of MtMce4A$_{39\text{-}140}$***

Purified MtMce4A$_{39\text{-}140}$ and SeMet labelled-MtMce4A$_{39\text{-}140}$ were concentrated up to 7.5 mg $ml^{-1}$ and used for all the crystallization experiments. Crystallization was performed using sitting drop vapor diffusion method at three different drop ratios (100:150, 150:150, 150:100 nl; protein: reservoir) at 22 °C. Crystals were observed in all three-drop ratios. Crystals were transferred to reservoir solution containing 25% of the appropriate cryoprotectant and flash-frozen in liquid nitrogen.

The data for both, the native MtMce4A$_{39\text{-}140}$ and SeMet-MtMce4A$_{39\text{-}140}$ crystals were collected at Biomax beamline, MaxIV (Lund, Sweden). The data processing and scaling was done using XDS (13) and AIMLESS (14), respectively, which suggested that the space group was $P6_1$ or $P6_5$.The SeMet-MtMce4A$_{39\text{-}140}$ structure was solved using SeMet-SAD phasing using the CRANK2 (15) pipeline. During hand-determination, the space group $P6_5$ was chosen based on figure of merit (FOM). The model obtained from CRANK2 pipeline was further build manually and the resulting R-free/R-work was 0.37 and 0.34 (with total 20 heavy atom sites used). This refined model generated from SeMet-MtMce4A$_{39\text{-}140}$ was used as the initial model for expert mode molecular replacement (Expert-MR) of PHASER (16) to determine the structure of native MtMce4A$_{39\text{-}140}$. The output from Expert-MR was used as an initial model for Autobuilding in Phenix (17). Further model building was done manually. Final refinement steps gave a R-free and R-work of 0.23 and 0.19 respectively. This final model of native-MtMce4A$_{39\text{-}140}$ was then again used to refine the SeMet-MtMce4A$_{39\text{-}140}$ data, which gave a final R-free and R-work of 0.24 and 0.21 respectively.

### Supplementary Results

#### *Recombinant expression and purification of MtMce1A-1F proteins*

Initially, the full length MtMce1A-1F were cloned and their expression in *E. coli* tested individually which showed that all the MtMce1A-1F individual proteins were successfully expressed. From these, MtMce1A, MtMce1B, MtMce1C and MtMce1F were selected for further purification trials as the expression for MtMce1D and MtMce1E was very low. Given that, all the Mce SBPs have N-terminal transmembrane domain, their solubility was assayed in buffers containing detergents such as DDM, Fos-choline-12 (FC-12) and Dodecyl nonaethylene glycol ether ($C_{12}E_9$) as mentioned in the methods section. Among these MtMce1A, MtMce1C, and MtMce1F were purified in the presence of DDM, whereas MtMce1B was purified only in the presence of FC-12. The major peak for the eluted proteins were between 10 and 12 ml in a 24 ml Superdex 200 10/300 column. An additional 8 ml peak was observed, suggesting the presence of soluble aggregates even in the presence of detergents.

Subsequently, the transmembrane domain deleted constructs were generated for MtMce1A and, MtMce1B (MtMce1A$_{38\text{-}454}$ and MtMce1B$_{29\text{-}346}$) to test if they could be purified without detergents. Surprisingly, even after deleting the transmembrane region, the proteins still required detergents for their purification (Fig. S2A). Also, as the expression of MtMce1E was very less, TM domain deleted construct was also made for MtMce1E (MtMce1E$_{37\text{-}390}$). Interestingly, the deletion resulted in enhanced expression of MtMce1E$_{37\text{-}390}$ when compared to the full length MtMce1E. However, for this construct a combination of DDM and $C_{12}E_9$ detergent was needed in different steps to purify it further (Supplementary Table 7). The TM domain deleted MceA-F SBPs almost eluted in the same volume as the full length MtMce1A-1F proteins. As the deletion of TM domain did not preclude the use of detergent, it was not generated for MtMce1C. However, in case of MtMce1F recombinant protein expression, heavy degradation was observed. Based on the mass spectrometry analysis of the bands corresponding to degradation as well as sequence analysis, we predicted that the degradation could mainly be at the extended tail domain. The tail domain of Mce1D is also long and has similarity with the tail domain of Mce1F. Therefore, transmembrane domain and tail domain deleted shorter constructs for MtMce1D (MtMce1D$_{44\text{-}314}$) and MtMce1F (MtMce1F$_{30\text{-}314}$) were generated and these constructs showed significantly less degradation (Fig. S2A). Intriguingly, even these constructs could be purified only in the presence of detergents.

#### *Recombinant expression and purification of MtMce4A-4F proteins*

In parallel, full-length MtMce4A-4F individual constructs were successfully generated in *E. coli.* Expression tests showed that the MtMce4F had the least expression. After initial detergent screening, MtMce4A, MtMce4C, MtMce4D and MtMce4F were purified in buffers containing the detergent DDM, whereas MtMce4B and MtMce4E required FC-12 for their purification. Gel filtration profile of MtMce4A, MtMce4C and MtMce4F showed that the aggregated protein peak (8 ml) was well separated from protein-detergent peak (~12 ml). In case of MtMce4B, MtMce4D and MtMce4E the aggregated protein and protein-detergent complex (PDC) eluted together in one broad peak. MtMce4B, MtMce4D and MtMce4F showed heavy degradation on SDS PAGE (Fig. 2B). As similar degradation was also observed for MtMce1F, it is possible that also in MtMce4B, MtMce4D and MtMce4F the degradation could be in the unstructured tail domain.

***Conformational changes in MtMce1A$_{36\text{-}148}$ and MtMce4A$_{39\text{-}140}$***

Comparison of the secondary structure content of MtMce4A$_{39\text{-}140}$ calculated from the CD spectrum with the crystal structure showed higher β sheet content (39%) in crystal than from the experimental CD spectra (28%) data (Table S3A). The result indicate that the protein is more structured in the crystallization condition and attain the MCE β-barrel fold. Interestingly, during thermal melting analysis when the temperature was gradually increased from 22 °C to 92 °C, a broad shift in the peak was observed between 220-240 nm corresponding to increase in secondary structures (Fig. S8B). This also aligned with the BeStSel analysis, which showed an increase in the β-sheet content with increasing temperatures. The β-sheet content at 72 °C was slightly higher (33%) indicating the unusual property of heat-induced conformational change of MtMce4A$_{39\text{-}140}$. However, the data after 72 °C was not reliable due to poor fitting of the data at 190-200 nm. Besides, the peak shift disappeared upon recooling of MtMce4A$_{39\text{-}140}$ indicating the temperature dependent reversible nature of this conformational change (Fig. S9B).

Similar thermal melting analysis of MtMce1A$_{36\text{-}148}$ showed conformational change upon heating. The initial peak at 198 nm, shifted to a broader range of 205-230 nm when the protein was heated from 22 °C to 92 °C (Fig. S8A). Moreover, the peaks at 210-230 nm were stable (not reversible) when the sample was recooled, whereas the peak between 205-210 nm disappeared during recooling (Fig. S9A). By visually looking at the spectra at 22 °C and 92 °C, one can interpret that MtMce1A also attains more secondary structure upon heating as observed in MtMce4A$_{39\text{-}140}$. Intriguingly, the deconvolution analysis (200-250 nm) of these peaks for MtMce1A$_{36\text{-}148}$ in both CDNN and BeStSel software suggested that only the spectra at 22 °C have higher β-sheet content and the β-sheet content reduced gradually upon heating. This overall indicates the challenges in the interpretation of CD spectra of β-sheet rich proteins even with the best available programs. In any case, we can clearly see that both MtMce1A$_{36\text{-}148}$ and MtMce4A$_{39\text{-}140}$ undergoes conformational change upon heating. It is possible that in the purified conditions, both MtMce1A$_{36\text{-}148}$ and MtMce4A$_{39\text{-}140}$ are in non-native conformations and MtMce4A$_{39\text{-}140}$ attains native conformation in the crystallization buffer.

**Supplementary Figures**

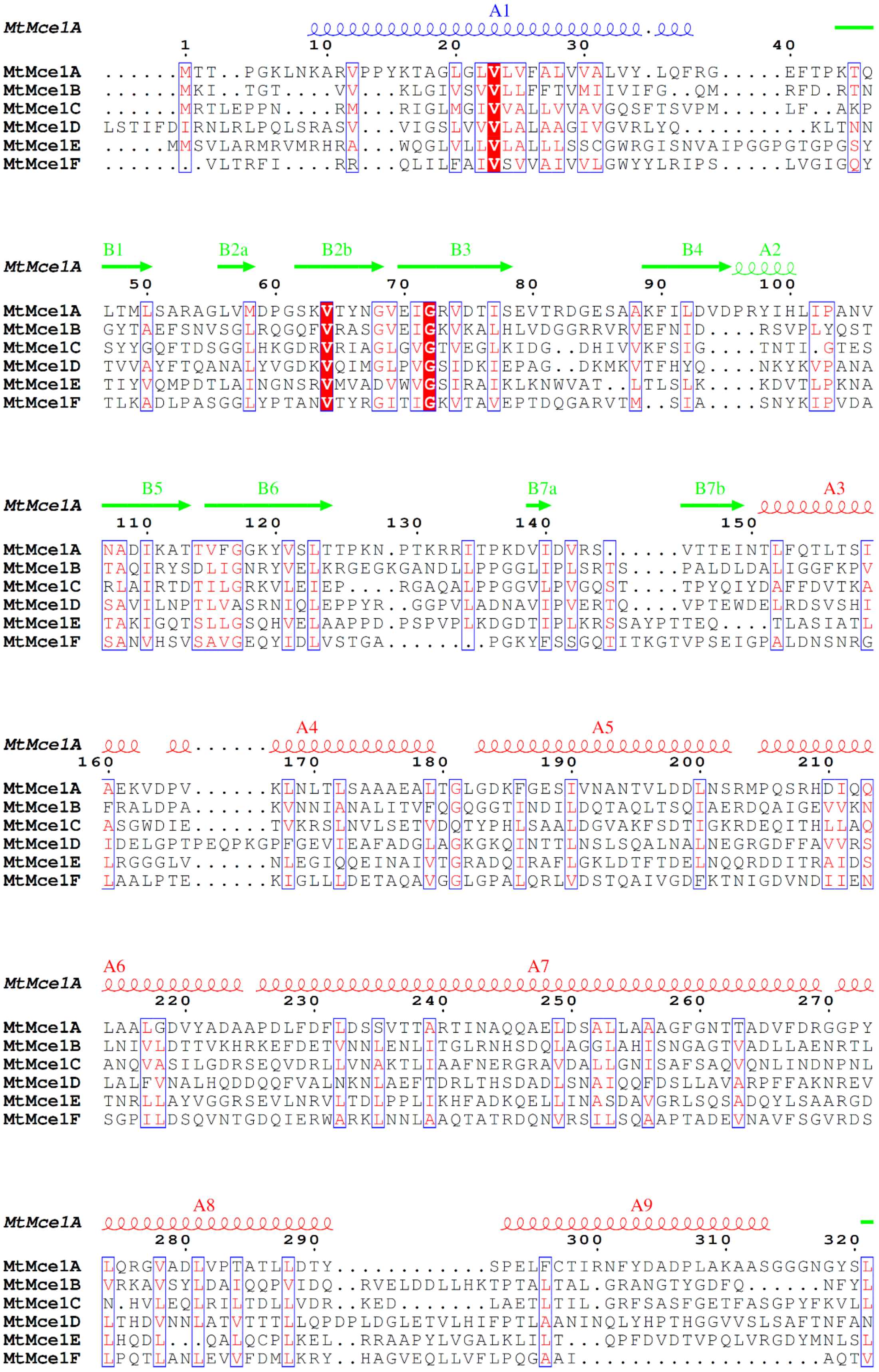

```
              B8   B9      A10     A11                               A12
MtMce1A
                  330       340          350         360
MtMce1A   RTNSEILSGIGISLLSPLALATN....GAAIGIGLVAG...LIAPPLAVAAN.......L
MtMce1B   .................CDL...............................QIK..W
MtMce1C   .................ANLVPG.................QILQPFVD....AAFK..K
MtMce1D   ................PMEFICSSIQAGSRLGYQESAELCAQYLAPVLDAIKFNYFPFGL
MtMce1E   ................TLDLTYS.................AIDNA........FLT..G
MtMce1F   .................LTPTPG.................AAQLPLAPAINYPPPC..L
```

```
                                                                  B10
MtMce1A
          370          380        390         400         410        420
MtMce1A   AGALPG...IVGGAPNPYTYPEN.LPRVNARGGPGGAPGCWQPITRDLWPAPYLVMDTGA
MtMce1B   NGFQAG......GPVRT....VKLFSQPTGRCTP...Q.....................
MtMce1C   RGISPE......DFWRSAGLPAYRWPDPNGTRFP...NGAPPP........PPPVLEGTP
MtMce1D   NVASTA.....STLPKEIAYSEPRLQPPNGYKDT.TVPGIWVP........DTPLSHR..
MtMce1E   TGFSGALRALEQSFGRDP...ETMIPD..IRYTP...NPNDAP........GGPLVERG.
MtMce1F   TGFLPA......SEWRSPADTSP.....................................
```

```
                                             A13
MtMce1A
              430                          440         450
MtMce1A   SLAPYNHMEVGSPY................AVEYVWGRQVGDNTIN.............
MtMce1B   ...........................................................
MtMce1C   E.HPGPAVPPGSPCSYTPPADGLPRPWDPLPCANLTQGPFGG.................
MtMce1D   ....................NTQPGWVVAPGMQGV...QVGPITQGLLTPESLAELM..
MtMce1E   ..........NRQC.............................................
MtMce1F   .....RPLPSGTYCKIPQDAQLQVRGARNIPCVDVLGKRAATPKEC..RSKDPYVPLGTN
```

```
MtMce1A

MtMce1A   ...........................................................
MtMce1B   ...........................................................
MtMce1C   PDFPAPLDVATSP..PNPDGPPPAPGLPIAGRPG......EVPPNVPGTPVPIPQE.APP
MtMce1D   ...............GGPDIAPPSSGLQTPPGPPN.........AYDEYPV.........
MtMce1E   ...........................................................
MtMce1F   PWFGDPNQILTCPAPGARCDQPVKPGLVIPAPSINTGLNPAPADQVQGTPPPVSDPLQRP
```

```
MtMce1A

MtMce1A   ...........................................................
MtMce1B   ...........................................................
MtMce1C   GARTL.PLG.PAPGPAPPPAAPGPPAPPGPGPQLPAPFINPGGTGGS...GVTGGSEN..
MtMce1D   ....LPPIGLQAPQVPIPPPPPGPDVIPGPVPPTPAPVGAPLPA......EAGGGQ....
MtMce1E   ...........................................................
MtMce1F   GSGTVQCNG.QQPNPCVYTPTSGPSAVYSPA...SGELVGPDGVKYAVANSSTTGDDGWK
```

```
MtMce1A

MtMce1A   .......
MtMce1B   .......
MtMce1C   .......
MtMce1D   .......
MtMce1E   .......
MtMce1F   EMLAPAS
```

**Supplementary Figure 1**: Multiple sequence alignment of MtMce1A-1F. The secondary structure elements of the MCE domain are based on the crystal structure of MtMce4A$_{39\text{-}140}$ and the remaining domains are from secondary structure prediction.

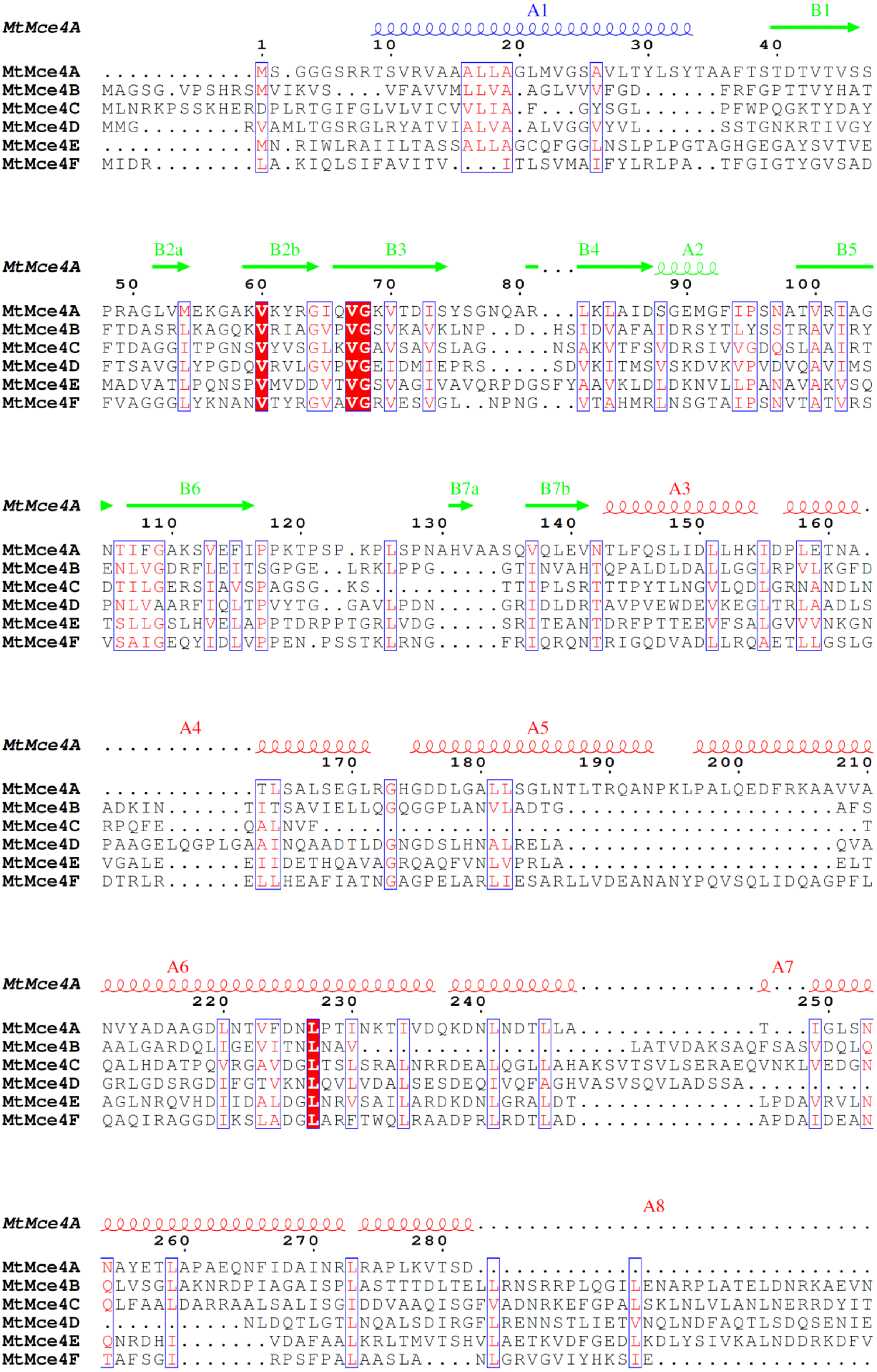
A1
B1
MtMce4A
1 10 20 30 40
MtMce4A ............MS.GGGSRRTSVRVAAALLAGLMVGSAVLTYLSYTAAFTSTDTVTVSS
MtMce4B MAGSG.VPSHRSMVIKVS....VFAVVMLLVA.AGLVVVFGD......FRFGPTTVYHAT
MtMce4C MLNRKPSSKHERDPLRTGIFGLVLVICVVLIA.F...GYSGL......PFWPQGKTYDAY
MtMce4D MMG........RVAMLTGSRGLRYATVIALVA.ALVGGVYVL......SSTGNKRTIVGY
MtMce4E ............MN.RIWLRAIILTASSALLAGCQFGGLNSLPLPGTAGHGEGAYSVTVE
MtMce4F MIDR........LA.KIQLSIFAVITV....ITLSVMAIFYLRLPA..TFGIGTYGVSAD
B2a B2b B3 B4 A2 B5
MtMce4A
50 60 70 80 90 100
MtMce4A PRAGLVMEKGAKVKYRGIQVGKVTDISYSGNQAR...LKLAIDSGEMGFIPSNATVRIAG
MtMce4B FTDASRLKAGQKVRIAGVPVGSVKAVKLNP..D..HSIDVAFAIDRSYTLYSSTRAVIRY
MtMce4C FTDAGGITPGNSVYVSGLKVGAVSAVSLAG.....NSAKVTFSVDRSIVVGDQSLAAIRT
MtMce4D FTSAVGLYPGDQVRVLGVPVGEIDMIEPRS.....SDVKITMSVSKDVKVPVDVQAVIMS
MtMce4E MADVATLPQNSPVMVDDVTVGSVAGIVAVQRPDGSFYAAVKLDLDKNVLLPANAVAKVSQ
MtMce4F FVAGGGLYKNANVTYRGVAVGRVESVGL..NPNG...VTAHMRLNSGTAIPSNVTATVRS
B6 B7a B7b A3
MtMce4A
110 120 130 140 150 160
MtMce4A NTIFGAKSVEFIPPKTPSP.KPLSPNAHVAASQVQLEVNTLFQSLIDLLHKIDPLETNA.
MtMce4B ENLVGDRFLEITSGPGE..LRKLPPG.....GTINVAHTQPALDLDALLGGLRPVLKGFD
MtMce4C DTILGERSIAVSPAGSG..KS..........TTIPLSRTTTPYTLNGVLQDLGRNANDLN
MtMce4D PNLVAARFIQLTPVYTG..GAVLPDN.....GRIDLDRTAVPVEWDEVKEGLTRLAADLS
MtMce4E TSLLGSLHVELAPPTDRPPTGRLVDG.....SRITEANTDRFPTTEEVFSALGVVVNKGN
MtMce4F VSAIGEQYIDLVPPEN.PSSTKLRNG.....FRIQRQNTRIGQDVADLLRQAETLLGSLG
A4 A5
MtMce4A
170 180 190 200 210
MtMce4A ............TLSALSEGLRGHGDDLGALLSGLNTLTRQANPKLPALQEDFRKAAVVA
MtMce4B ADKIN......TITSAVIELLQGQGGPLANVLADTG.....................AFS
MtMce4C RPQFE......QALNVF..........................................T
MtMce4D PAAGELQGPLGAAINQAADTLDGNGDSLHNALRELA.....................QVA
MtMce4E VGALE......EIIDETHQAVAGRQAQFVNLVPRLA.....................ELT
MtMce4F DTRLR......ELLHEAFIATNGAGPELARLIESARLLVDEANANYPQVSQLIDQAGPFL
A6 A7
MtMce4A
220 230 240 250
MtMce4A NVYADAAGDLNTVFDNLPTINKTIVDQKDNLNDTLLA..............T...IGLSN
MtMce4B AALGARDQLIGEVITNLNAV.....................LATVDAKSAQFSASVDQLQ
MtMce4C QALHDATPQVRGAVDGLTSLSRALNRRDEALQGLLAHAKSVTSVLSERAEQVNKLVEDGN
MtMce4D GRLGDSRGDIFGTVKNLQVLVDALSESDEQIVQFAGHVASVSQVLADSSA..........
MtMce4E AGLNRQVHDIIDALDGLNRVSAILARDKDNLGRALDT..............LPDAVRVLN
MtMce4F QAQIRAGGDIKSLADGLARFTWQLRAADPRLRDTLAD..............APDAIDEAN
A8
MtMce4A
260 270 280
MtMce4A NAYETLAPAEQNFIDAINRLRAPLKVTSD...............................
MtMce4B QLVSGLAKNRDPIAGAISPLASTTTDLTELLRNSRRPLQGILENARPLATELDNRKAEVN
MtMce4C QLFAALDARRAALSALISGIDDVAAQISGFVADNRKEFGPALSKLNLVLANLNERRDYIT
MtMce4D ...........NLDQTLGTLNQALSDIRGFLRENNSTLIETVNQLNDFAQTLSDQSENIE
MtMce4E QNRDHI.......VDAFAALKRLTMVTSHVLAETKVDFGEDLKDLYSIVKALNDDRKDFV
MtMce4F TAFSGI.......RPSFPALAASLA....NLGRVGVIYHKSIE.................

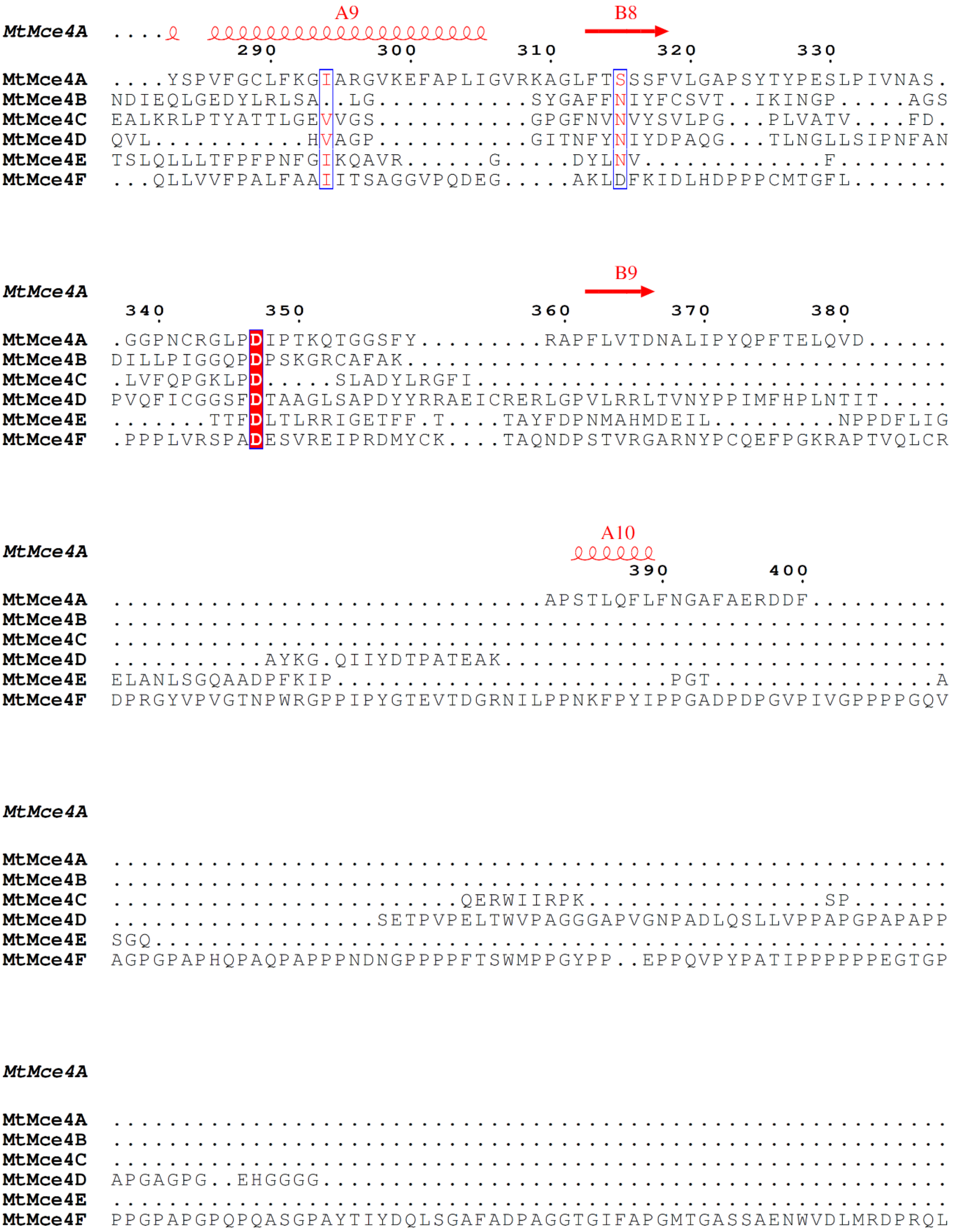


**Supplementary Figure 2**: Multiple sequence alignment of MtMce4A-4F. The secondary structure elements of the MCE domain are from the crystal structure of $MtMce4A_{39-140}$ and the remaining domains are from secondary structure prediction.

(A)

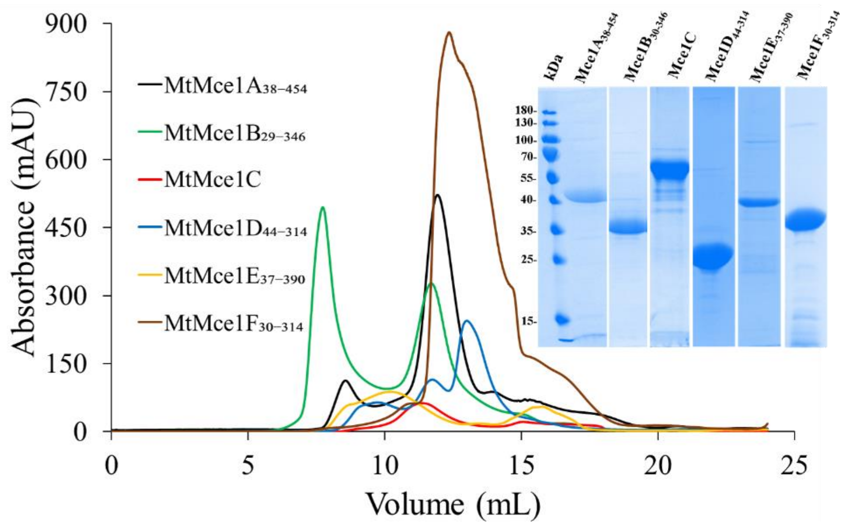


(B)

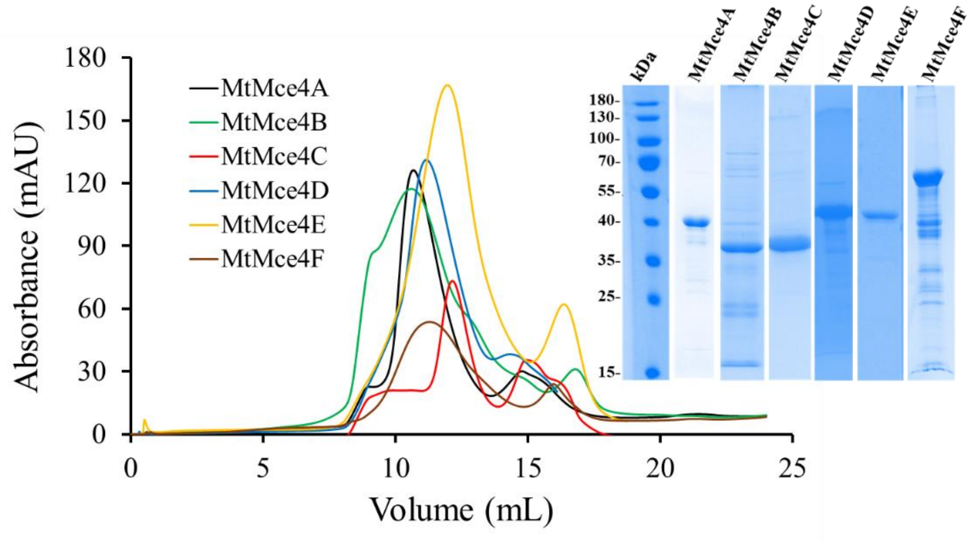


**Supplementary Figure 3**: SEC elution profiles of **(A)** selected individual MtMce1A-1F SBPs and **(B)** individual MtMce4A-4F SBPs on a 24 ml Superdex 200 10/300 column. The protein samples were analyzed on a 12 % SDS-PAGE (inset).

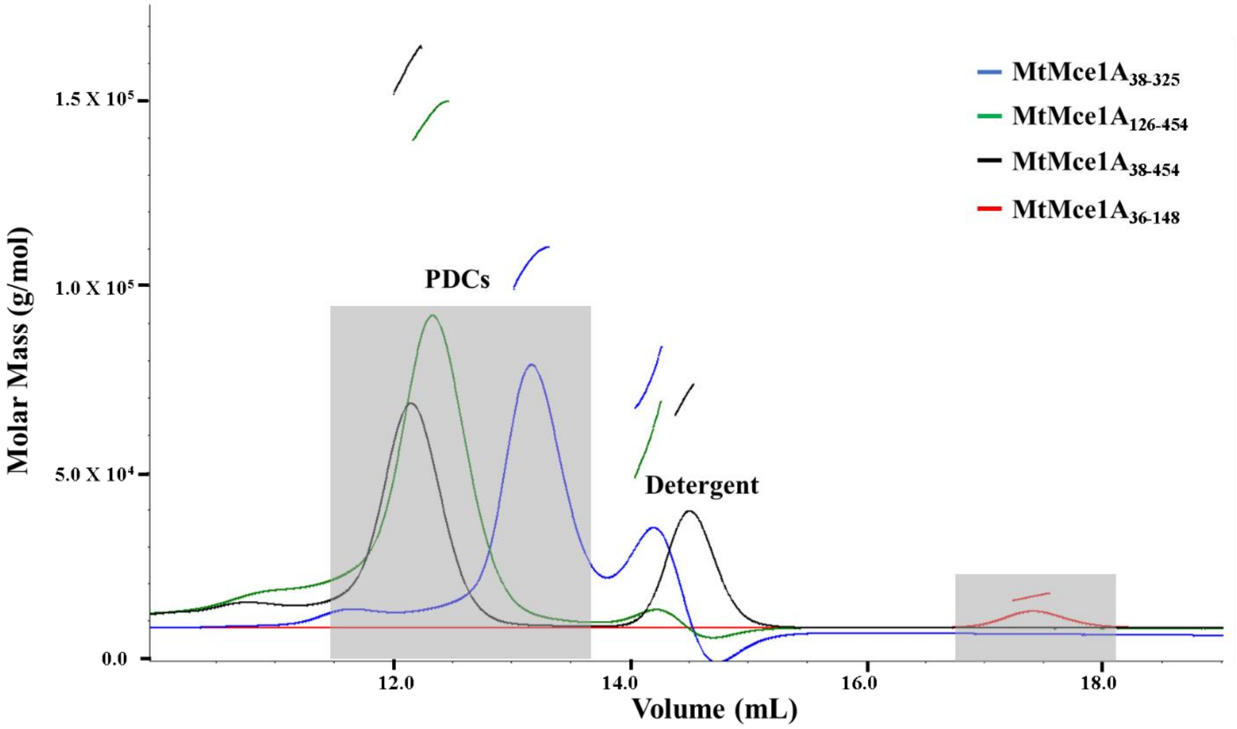


**Supplementary Figure 4**: SEC-MALS profile of Mce1A$_{36-148}$ (red), Mce1A$_{38-325}$ (blue), Mce1A$_{126-454}$ (green), and Mce1A$_{38-454}$ (black). Mce1A$_{36-148}$ has a single scattering peak at ~17.5 ml. Whereas other Mce1A domains are purified in DDM showed two scattering peaks corresponds to protein-detergent complex (12-14 ml) and empty detergent micelle. All the samples were injected on a 24 ml Superdex 200 10/300 column.

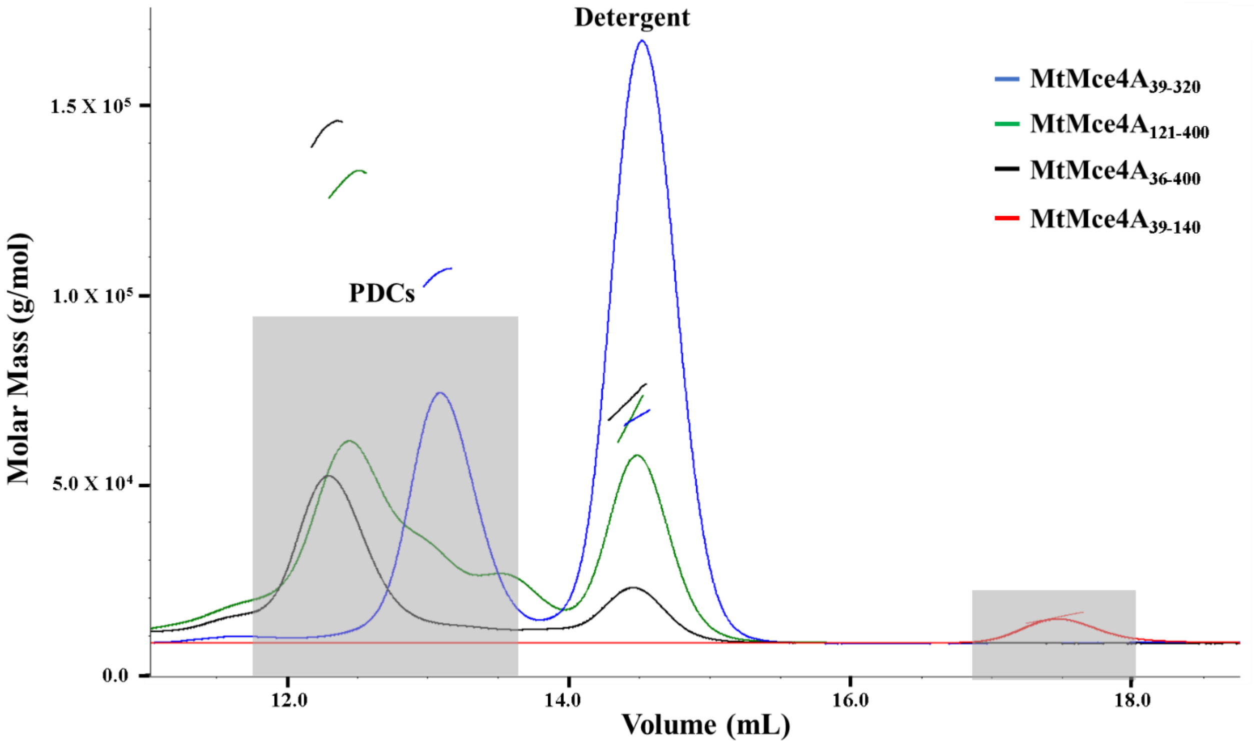


**Supplementary Figure 5**: SEC-MALS profile of Mce4A$_{39-140}$ (red), Mce4A$_{39-320}$ (blue), Mce4A$_{121-400}$ (green), and Mce4A$_{36-400}$ (black). Mce4A$_{39-140}$ has a single scattering peak at ~17.5 ml. Whereas other Mce4A domains are purified in DDM showed two scattering peaks corresponds to protein-detergent complex (12-14 ml) and empty detergent micelle. All the samples were injected on a 24 ml Superdex 200 10/300 column.

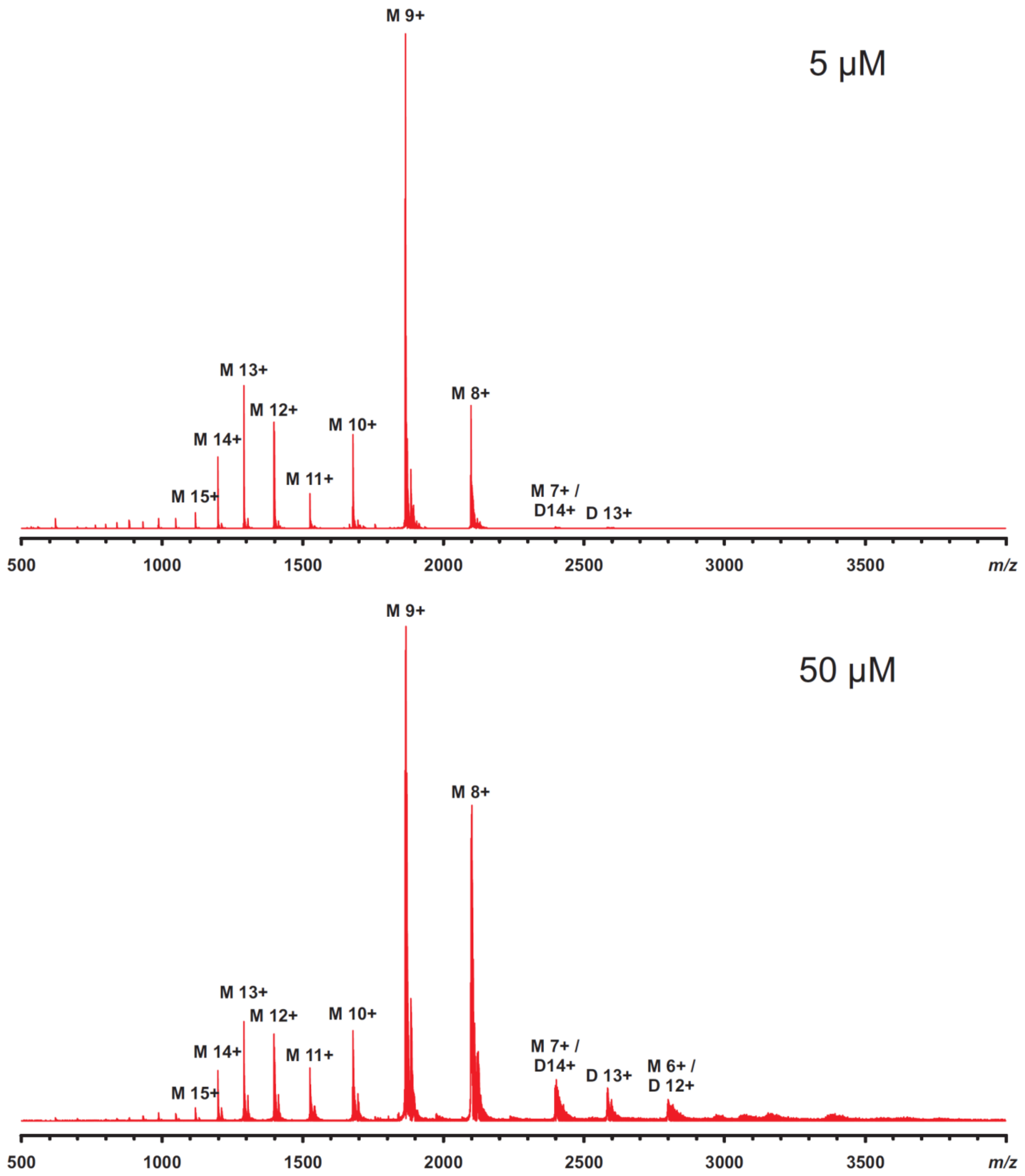


**Supplementary Figure 6**: Native mass spectra of MtMce1A$_{36\text{-}148}$ at 5 µM and 50 µM concentration in 20 mM ammonium acetate buffer, pH 6.8.

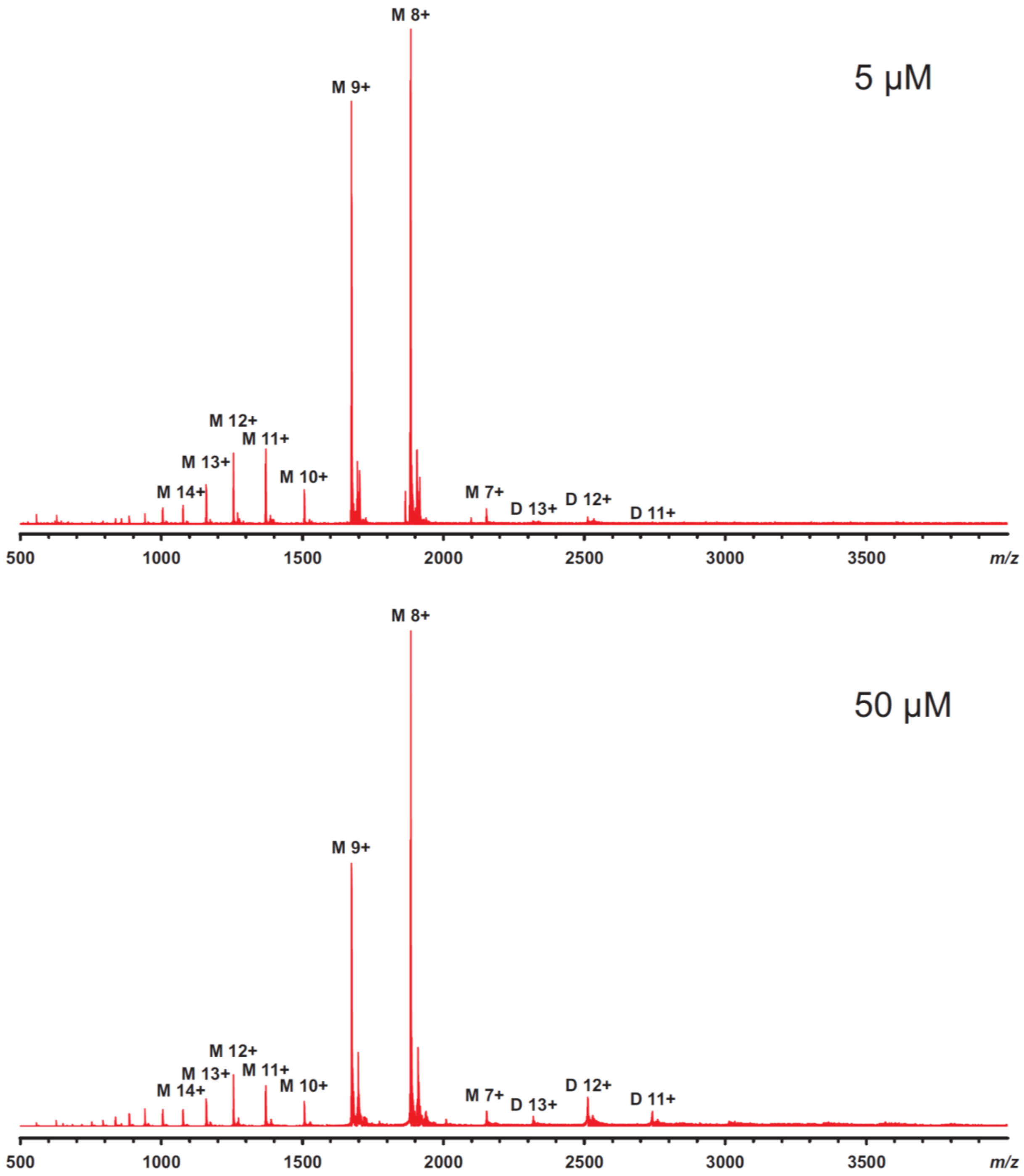


**Supplementary Figure 7**: Native mass spectra of MtMce4A$_{39\text{-}140}$ at 5 µM and 50 µM concentration in 20 mM ammonium acetate buffer, pH 6.8.

(A)

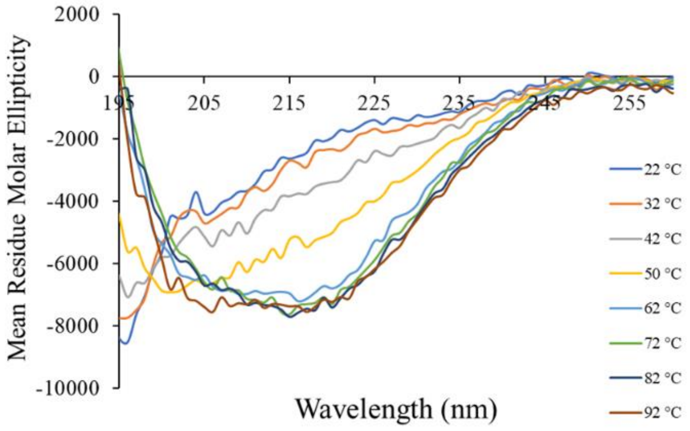


(B)

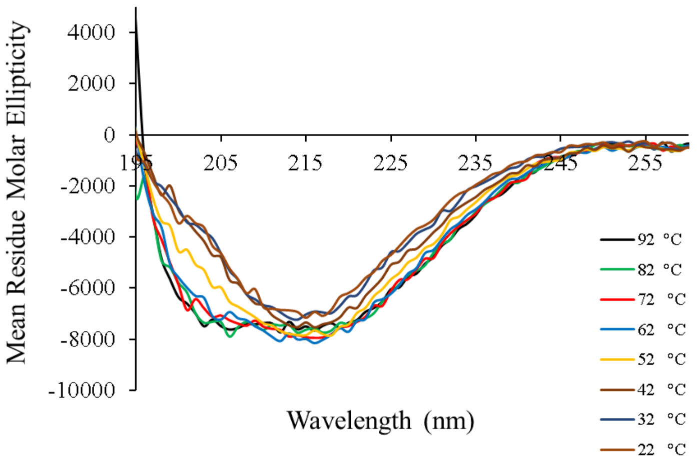


**Supplementary Figure 8: (A)** CD spectra of MtMce1A$_{36-148}$ from 22 °C to 92 °C. **(B)** CD spectra of MtMce1A$_{36-148}$ from 92 °C to 22 °C.

(A)

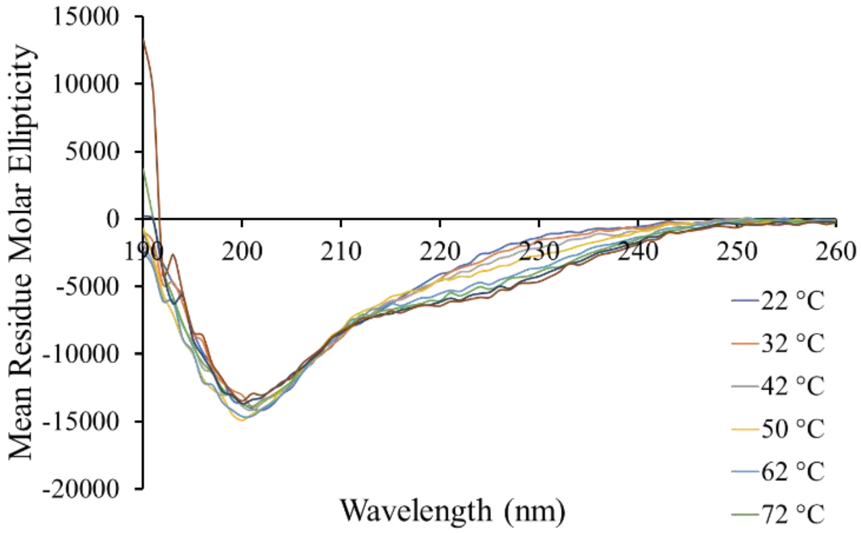


(B)

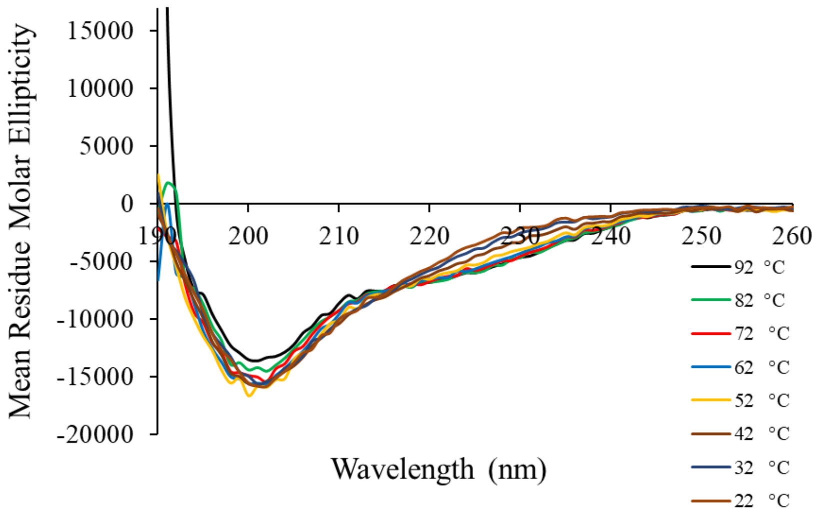


**Supplementary Figure 9: (A)** CD spectra of MtMce4A$_{39\text{-}140}$ from 22 °C to 72 °C. **(B)** CD spectra of MtMce4A$_{39\text{-}140}$ from 92 °C to 22 °C.

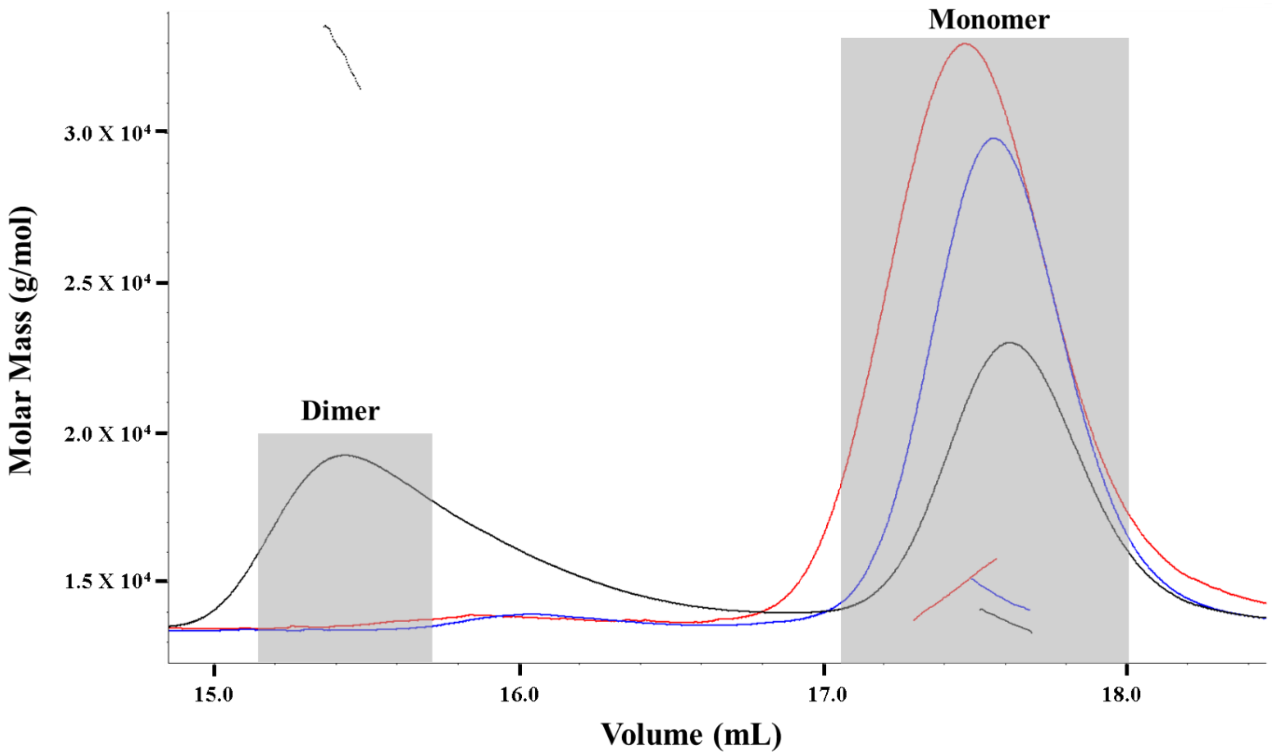


**Supplementary Figure 10**: Comparison of SEC-MALS of Mce4A$_{39\text{-}140}$ in three different conditions: Mce4A$_{39\text{-}140}$ in purification buffer (50 mM MOPS, 350 mM NaCl, 10% Glycerol, pH 7.0) in blue, Mce4A$_{39\text{-}140}$ in crystallization buffer (0.1 M MES; 0.7 M ammonium sulfate, pH 6.0) in red, and Mce4A$_{39\text{-}140}$ in crystallization buffer heated up to 50°C in black.

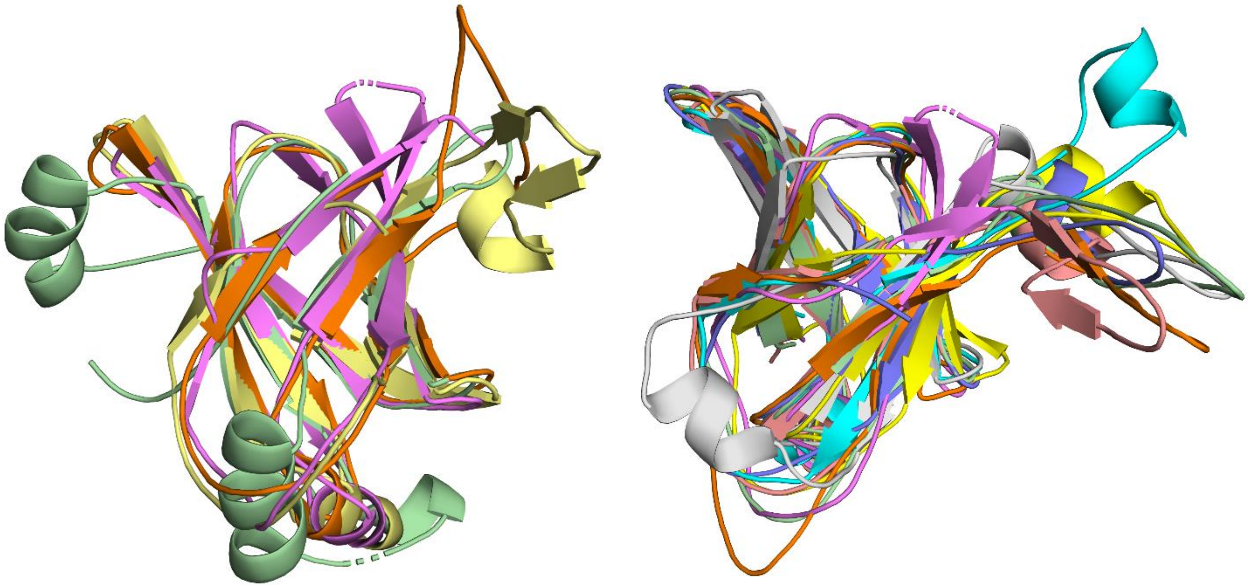

**Supplementary Figure 11**: **(A)** Structural superposition of MtMce4A$_{39-140}$ (pink) with MCE domain of EcPqiB$_{1}$ (orange) EcPqiB$_{2}$ (yellow), EcPqiB$_{3}$ (green) monomers. **(B)** Structural overlap of MtMce4A$_{39-140}$ (pink) with MCE domain of EcLetB$_{1}$(cyan), EcLetB$_{2}$(green), EcLetB$_{3}$(yellow), EcLetB$_{4}$(peach), EcLetB$_{5}$(gray), EcLetB$_{6}$(purple), and EcLetB$_{7}$(orange) monomers.

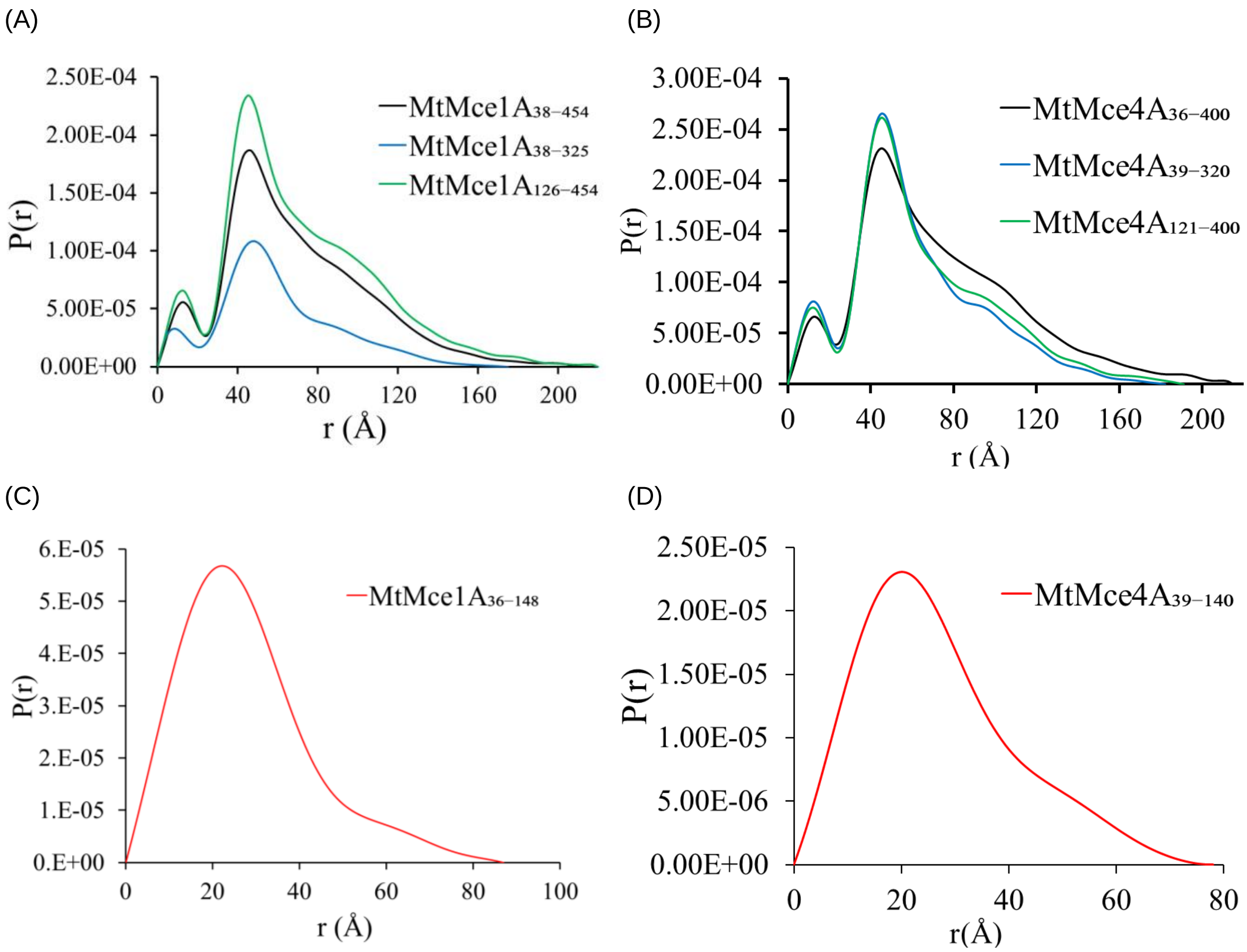


**Supplementary Figure 12**: **(A)** The P(r) curves of MtMce1A$_{38\text{-}454}$, MtMce1A$_{38\text{-}325}$ and MtMce1A$_{126\text{-}454}$. **(B)**The P(r) curves of MtMce4A$_{36\text{-}400}$, MtMce4A$_{39\text{-}320}$ and MtMce4A$_{121\text{-}400}$. **(C)** The P(r) curve of MtMce1A$_{36\text{-}148}$ and **(D)** MtMce4A$_{39\text{-}140}$.

(A) MtMce1A38-325

Extended Coiled-coil

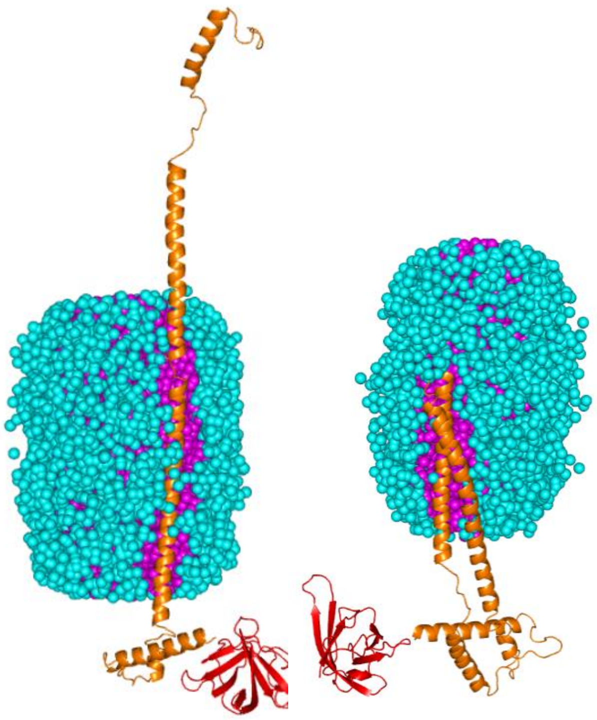

(B)

Extended

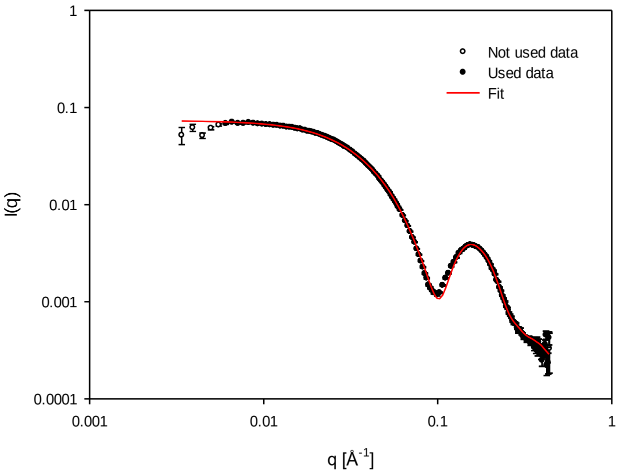


Coiled-coil

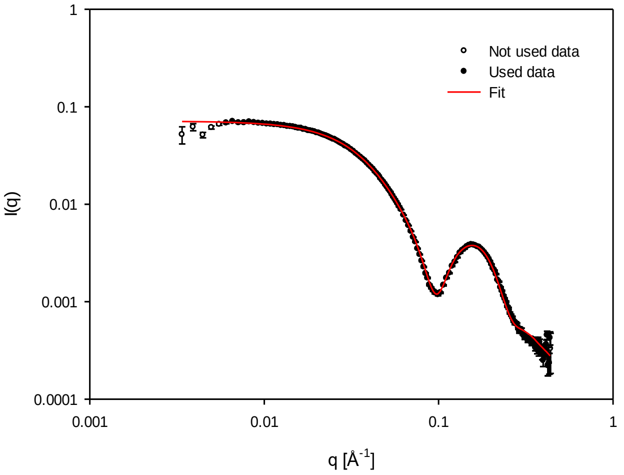


**Supplementary Figure 13**: **(A)** The extended (left) and coiled-coil (right) model of MtMce1A$_{38\text{-}325}$. **(B)** The fit of experimental SAXS data with the proposed models of MtMce1A$_{38\text{-}325}$.

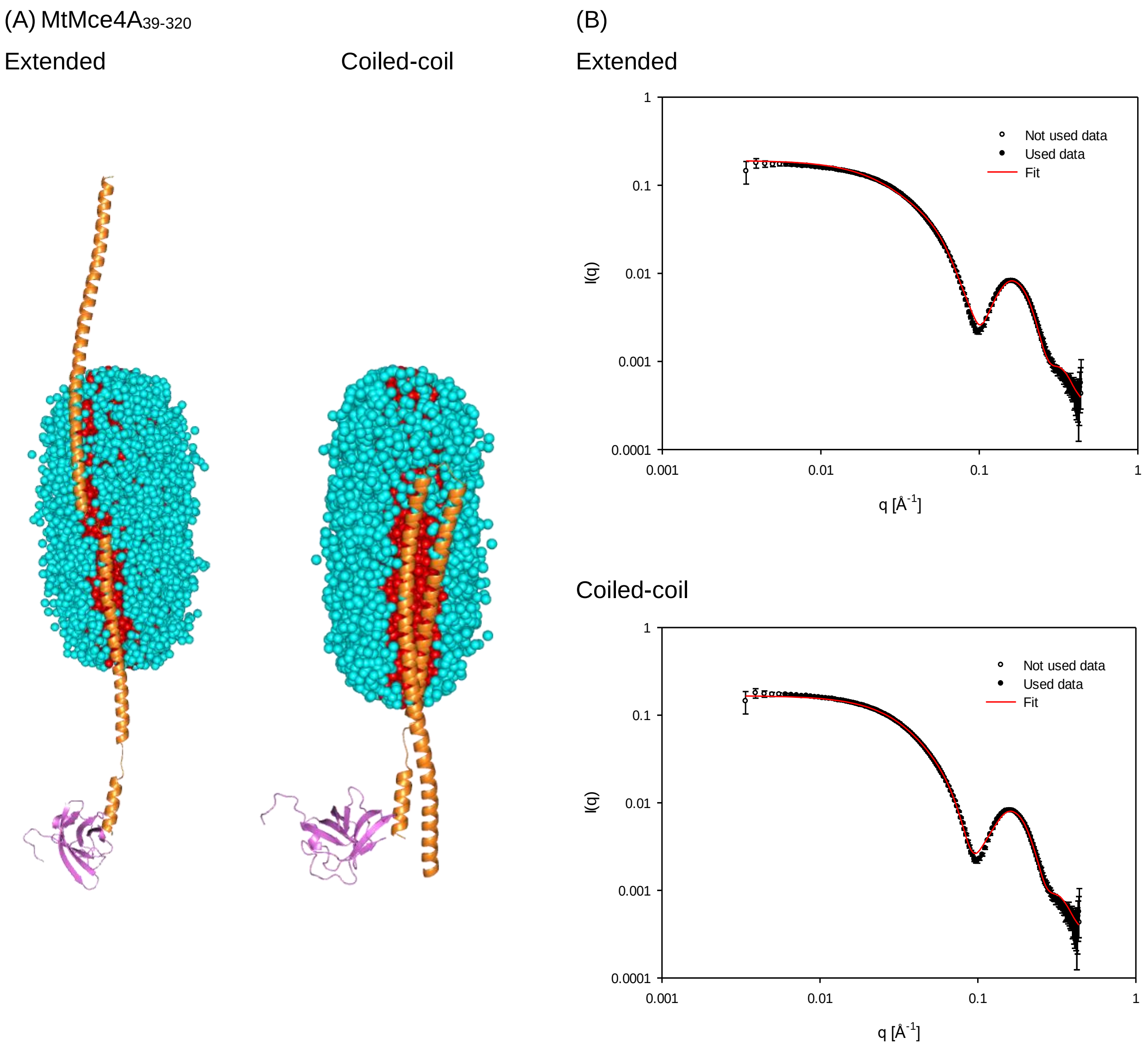


**Supplementary Figure 14**: **(A)** The extended (left) and coiled-coil (right) model of MtMce4A$_{39\text{-}320}$**. (B)** The fit of experimental SAXS data with the proposed models of MtMce4A$_{39\text{-}320}$.

(A) MtMce1A38-454

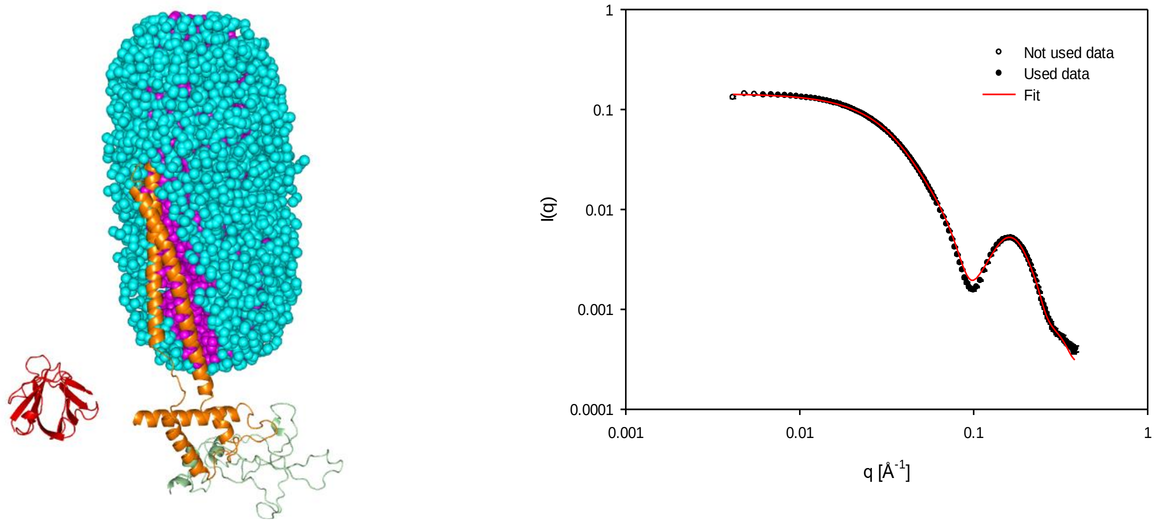


(B) MtMce4A36-400

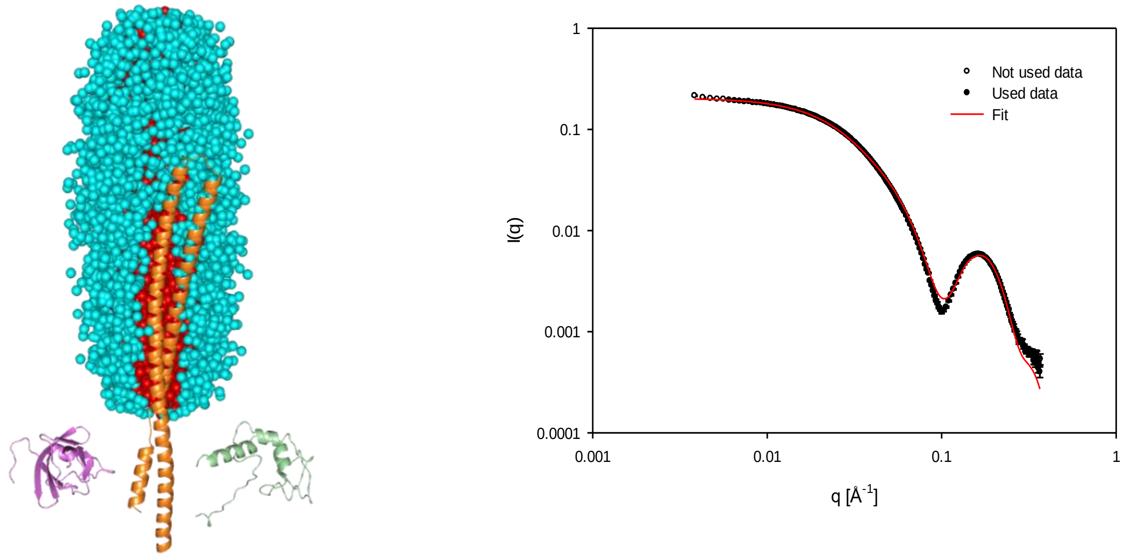


**Supplementary Figure 15**: **(A)** The coiled-coil model and the fit of experimental SAXS data with the proposed models of MtMce1A$_{38\text{-}454}$. **(B)** coiled-coil model and the fit of experimental SAXS data with the proposed model of MtMce4A$_{36\text{-}400}$.

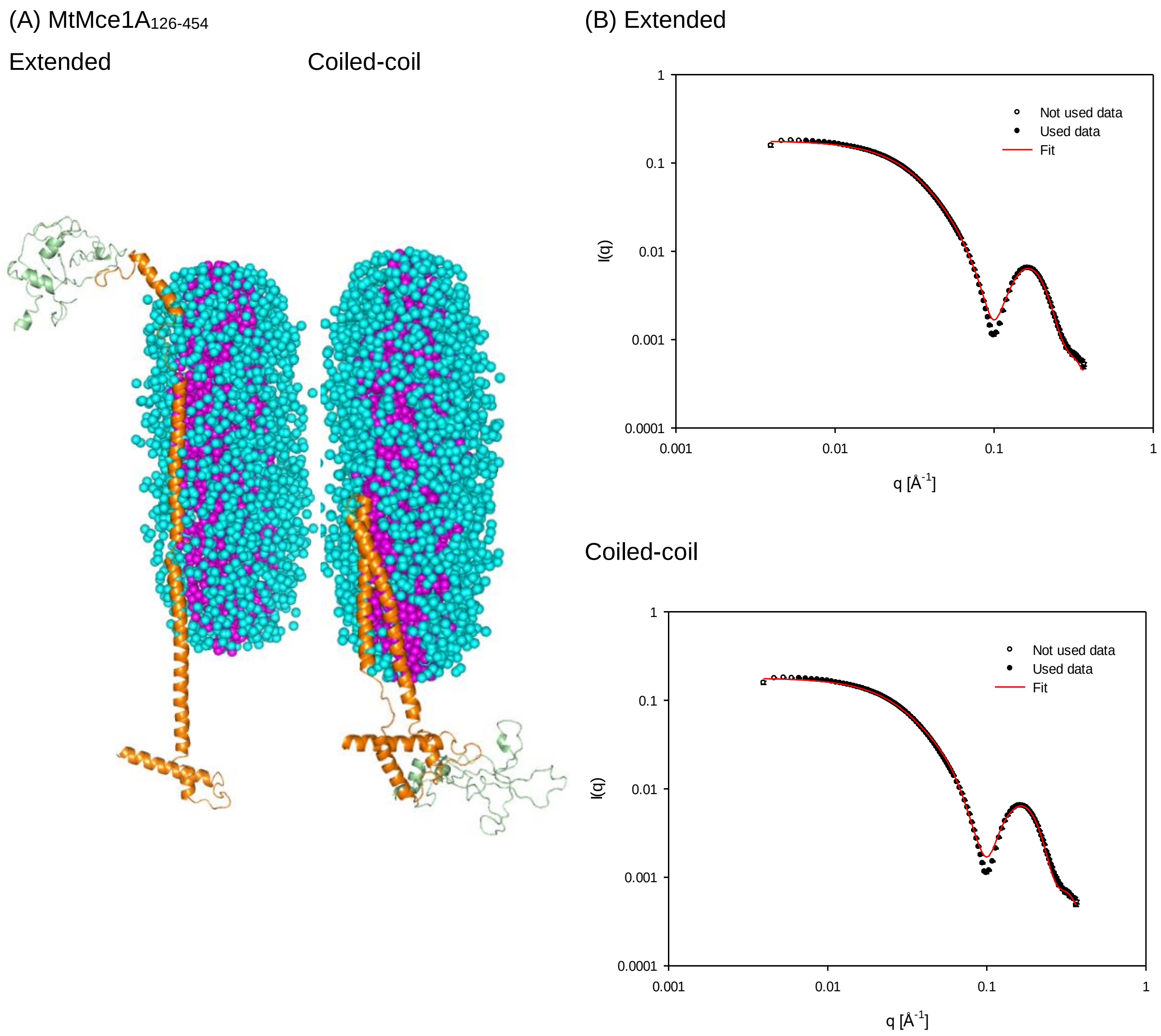


**Supplementary Figure 16**: **(A)** The extended and coiled-coil models of $MtMce1A_{126\text{-}454}$. **(B)** The fit of experimental SAXS data with the proposed models of $MtMce1A_{126\text{-}454}$.

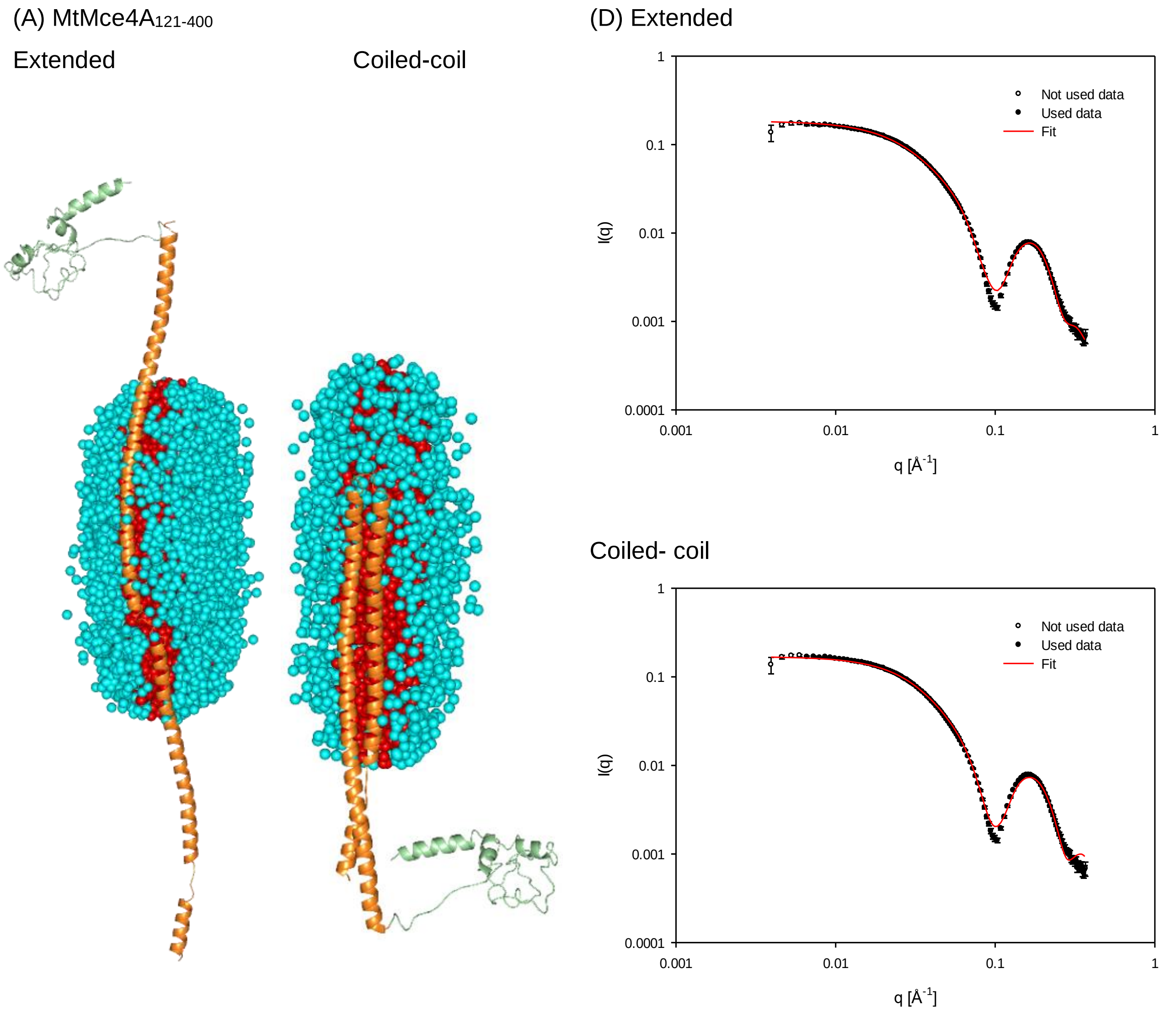


**Supplementary Figure 17**: **(A)** The extended and coiled-coil models of MtMce4A$_{121-400}$. **(B)** The fit of experimental SAXS data with the proposed models of MtMce4A$_{121-400}$.

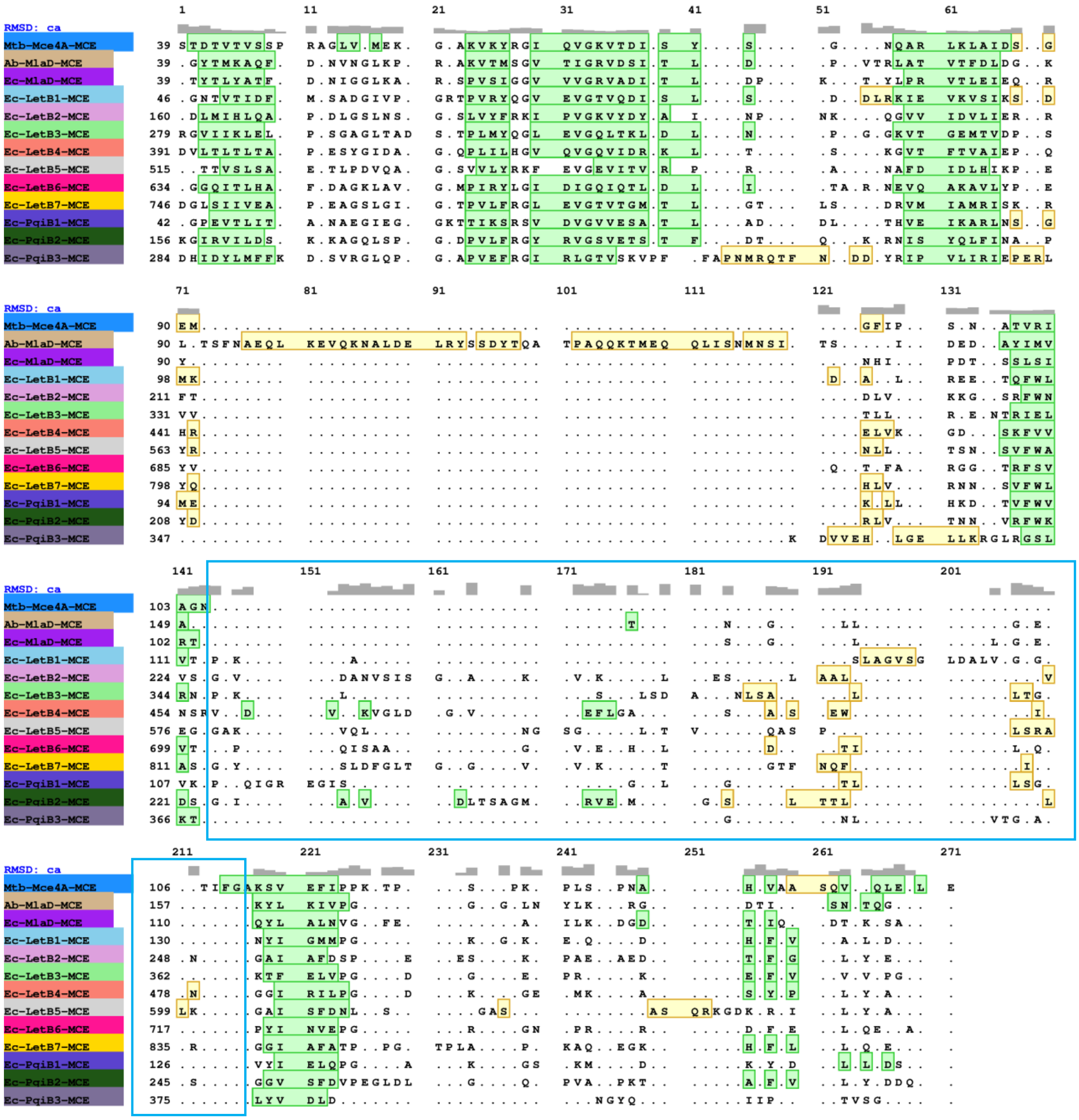

**Supplementary Figure 18**: The structure-based sequence aligment of all the known Mce SBP structures (EcMlaD, AbMlaD, EcPqiB1-3, and EcLetB1-7) with the MtMce4A39-140. The alignment was generated using matchmaker (chimera). The β-strands and α-helices are highlighted in green and yellow, respectively. The PLL loop is highlighted in blue box.

**Supplementary Tables**

**Table S1: Percent identity matrix for MtMce1A-1F created by Clustal Omega**

| **MtMce1A-1F** | | | | | | |
|---|---|---|---|---|---|---|
| | MtMce1A | MtMce1B | MtMce1C | MtMce1D | MtMce1E | MtMce1F |
| MtMce1A | 100.00 | 17.07 | 18.11 | 20.15 | 16.76 | 18.88 |
| MtMce1B | 17.07 | 100.00 | 21.52 | 18.64 | 17.77 | 16.13 |
| MtMce1C | 18.11 | 21.52 | 100.00 | 20.67 | 20.68 | 20.31 |
| MtMce1D | 20.15 | 18.64 | 20.67 | 100.00 | 17.63 | 18.41 |
| MtMce1E | 16.76 | 17.77 | 20.68 | 17.63 | 100.00 | 18.32 |
| MtMce1F | 18.88 | 16.13 | 20.31 | 18.41 | 18.32 | 100.00 |

**Table S2: Percent identity matrix for MtMce4A-4F created by Clustal Omega**

| **MtMce4A-4F** | | | | | | |
|---|---|---|---|---|---|---|
| | MtMce4A | MtMce4B | MtMce4C | MtMce4D | MtMce4E | MtMce4F |
| MtMce4A | 100.00 | 17.27 | 15.36 | 19.27 | 18.69 | 21.43 |
| MtMce4B | 17.27 | 100.00 | 19.55 | 22.65 | 16.16 | 17.54 |
| MtMce4C | 15.36 | 19.55 | 100.00 | 28.39 | 19.86 | 16.38 |
| MtMce4D | 19.27 | 22.65 | 28.39 | 100.00 | 17.03 | 20.11 |
| MtMce4E | 18.69 | 16.16 | 19.86 | 17.03 | 100.00 | 24.14 |
| MtMce4F | 21.43 | 17.54 | 16.38 | 20.11 | 24.14 | 100.00 |

**Table S3: Secondary structure content of MtMce1A and MtMce4A domains**

| | **Helix** | **Anti-Parallel β-strand** | **Parallel β-strand** | **β-turn** | **Random coil** |
|---|---|---|---|---|---|
| MtMce1A$_{36-148}$ | 2.3% | 40.6% | 3.7% | 13.5% | 39.9% |
| MtMce1A$_{38-325}$ | 27.9% | 13.3% | 9.8% | 18.0% | 33.7% |
| MtMce1A$_{126-454}$ | 30.6% | 11.7% | 8.8% | 17.6% | 30.6% |
| MtMce1A$_{38-454}$ | 38.3% | 8.5% | 7.2% | 16.3% | 25.7% |
| MtMce4A$_{39-140}$ | 6.7% | 24.2% | 3.9% | 13.2% | 52% |
| MtMce4A$_{39-320}$ | 28.0% | 12.8% | 9.9% | 17.9% | 34.8% |
| MtMce4A$_{121-400}$ | 33.8% | 10.5% | 7.8% | 17.1% | 26.8% |
| MtMce4A$_{36-400}$ | 37.5% | 8.3% | 7.1% | 16.3% | 25.6% |

**Table S4: Secondary structure comparison of $MtMce4A_{39-140}$ in solution and crystal structure**

| | In solution | Crystal structure |
|---|---|---|
| alpha-Helix (distorted) | 6.70 | 4.10 |
| β-strands (parallel and Anti-parallel) | 28.10 | 39.04 |
| β-turn | 13.20 | 8.90 |
| Others | 52.00 | 47.95 |

**Table S5: SAXS structural parameters for MtMce1A and MtMce4A soluble domains ($MtMce1A_{36-148}$, and $MtMce4A_{39-140}$)**

| | **$MtMce1A_{36-148}$** | **$MtMce4A_{39-140}$** |
|---|---|---|
| Beamline | B21, DLS, UK | B21, DLS, UK |
| Beam size (µm) | 250x250 | 250x250 |
| Detector | Pilatus 2M | Pilatus 2M |
| Wavelength (Å) | 1 | 1 |
| Camera length (m) | 4.014 | 4.014 |
| Protein concentration (mg $ml^{-1}$) | 2.0 | 1 |
| Volume injected (µl) | 30 | 30 |
| Buffer used | 50mM Tris, 350 mM NaCl, 10% Glycerol, pH 8.5 | 50mM MOPS, 350 mM NaCl, 10% Glycerol, pH 7.0 |
| Buffer subtraction | SCATTER | SCATTER |
| Primary analysis | PRIMUS | PRIMUS |
| *Ab-initio* shape | DAMMINN | DAMMINN |
| Atomic structure modelling | Robetta | Crystal structure+ Robetta |
| 3-D representation | PyMOL | PyMOL |
| **Guinier and P(r) analysis** | | |
| I(0) $cm^{-1}$ | 0.023 | 0.01 |
| $R_g$ (Å) | 21.2 | 20.76 |
| q range (Å) | | |
| $qR_g$ max | 1.29 | 1.29 |
| $D_{max}$ (A) | 87 | 80 |
| $\chi^2$ (total estimate from GNOM) | 0.59 | 0.71 |
| Porod volume, Po | 28759 | 22236 |
| Molecular mass derived from Qp, MoW, Vc, size & shape (kDa) | 11.13, 11.60, 15.44, 19.9 | 10.03, 10,30, 14.22, 16.37 |
| **DAMMIN parameters** | | |
| Symmetry, anisotropy assumption | P1 unknown | P1 unknown |
| $\chi^2$ | 1.26 | 1.104 |

| **Atomistic modelling** | | |
|---|---|---|
| Model used | Elongated and compact | Elongated and compact |
| **Fit** | | |
| $\chi^2$ | 4.2, 14.0 | 2.0, 10.0 |
| **SASDB accession codes** | SASDJU9 | SASDJV9 |

**Table S6: SAXS structural parameters for MtMce1A domains (MtMce1A$_{38-454}$, MtMce1A$_{126-454}$, and MtMce1A$_{38-325}$)**

| | **MtMce1A$_{38-454}$** | **MtMce1A$_{126-454}$** | **MtMce1A$_{38-325}$** |
|---|---|---|---|
| Beamline | B21, DLS, UK | B21, DLS, UK | B21, DLS, UK |
| Beam size (µm) | 250x250 | 250x250 | 250x250 |
| Detector | Pilatus 2M | Pilatus 2M | Pilatus 2M |
| Wavelength (Å) | 1 | 1 | 1 |
| Camera length (m) | 4.014 | 4.014 | 4.014 |
| SEC-SAXS column | Superdex 200 increase 3.2/300 | Superdex 200 increase 3.2/300 | Superdex 200 increase 3.2/300 |
| Protein concentration (mg ml$^{-1}$) | 5 | 5.2 | 6.0 |
| Volume injected (µl) | 55 | 55 | 55 |
| Flow rate (ml min$^{-1}$) | 0.075 | 0.15 | 0.06 |
| Buffer used | 50 mM Tris, 500 mM NaCl, 10% Glycerol, 5mM DDM, 1 mM β-ME, pH 8.0 | 50mM Tris, 350 mM Nacl, 10%glycerol, 5 mM DDM, 1mM β -ME, pH 7.5 | 50mM Tris, 500mM NaCl, 10% Glycerol, 5mM DDM, 1mM β -ME, pH 7.5 |
| Buffer subtraction | SCATTER | SCATTER | SCATTER |
| Primary analysis | PRIMUS | PRIMUS | PRIMUS |
| *Ab-initio* shape | - | - | - |
| Atomic structure modelling | Swiss-model, iTasser | Swiss-model, iTasser | Swiss-model, iTasser |
| 3-D representation | PyMOL | PyMOL | PyMOL |
| **Guinier and P(r) analysis** | | | |
| I(0) cm$^{-1}$ | 0.15 | 0.18 | 0.07 |
| $R_g$ (Å) | 52.8 | 54.86 | 45.82 |
| $qR_g$ max | 1.29 | 1.29 | 1.29 |
| $D_{max}$ (Å) | 216 | 220 | 175 |
| $\chi^2$ (total estimate from GNOM) | 0.63 | 0.62 | 0.67 |
| Porod volume, Po | 362264 | 440058 | 151292 |
| **R factor analysis** | | | |

| | | | |
|---|---|---|---|
| R factor $= \frac{\sum \lvert I(q_i) - I_{model}(q_i) \rvert}{\sum I(q_i)}$ (Extended model, coiled-coil) | 1.2 %, 1.3 % | 1.5 %, 2.2 % | 1.0 %, 0.9 % |
| Weighted R factor= $\left(\frac{\sum (I(q_i) - I_{model}(q_i))^2/\sigma(I(q_i))^2}{\sum I(q_i)^2/\sigma(I(q_i))^2}\right)^{0.5}$ (Extended model, coiled-coil) | 1.9 %, 2.4 % | 2.3 %, 3.3 % | 1.9 %, 1.6 % |
| **SASDB accession codes** | SASDK32 | SASDK22 | SASDJZ9 |

**Table S7: SAXS structural parameters for MtMce4A domains (MtMce4A$_{36-400}$, MtMce4A$_{121-400}$, and MtMce4A$_{39-320}$)**

| | **MtMce4A$_{36-400}$** | **MtMce4A$_{121-400}$** | **MtMce4A$_{39-320}$** |
|---|---|---|---|
| Beamline | B21, DLS, UK | B21, DLS, UK | B21, DLS, UK |
| Beam size (µm) | 250x250 | 250x250 | 250x250 |
| Detector | Pilatus 2M | Pilatus 2M | Pilatus 2M |
| Wavelength (Å) | 1 | 1 | 1 |
| Camera length (m) | 4.014 | 4.014 | 4.014 |
| SEC-SAXS column | Superdex 200 increase 3.2/300 | Superdex 200 increase 3.2/300 | Superdex 200 increase 3.2/300 |
| Protein concentration (mg ml$^{-1}$) | 5 | 5 | 5 |
| Volume injected (µl) | 55 | 55 | 55 |
| Flow rate (mL min$^{-1}$) | 0.075 | 0.15 | 0.06 |
| Buffer used | 50 mM Tris, 500 mM NaCl, 10% Glycerol, 5mM DDM, 1 mM β-ME, pH 8.0 | 50 mM Tris, 500 mM NaCl, 10% Glycerol, 5mM DDM, 1 mM β-ME, pH 8.0 | 50 mM Tris, 500 mM NaCl, 10% Glycerol, 5mM DDM, 1 mM β-ME, pH 8.5 |
| Buffer subtraction | SCATTER | SCATTER | SCATTER |
| Primary analysis | PRIMUS | PRIMUS | PRIMUS |
| *Ab-initio* shape | - | - | - |
| Atomic structure modelling | Mce4A$_{39-140}$ Crystal structure, iTasser | Mce4A$_{39-140}$ Crystal structure, iTasser | Mce4A$_{39-140}$ Crystal structure, iTasser |
| 3-D representation | PyMOL | PyMOL | PyMOL |
| **Guinier and P(r) analysis** | | | |
| I(0) cm$^{-1}$ | 0.20 | 0.17 | 0.17 |
| $R_g$ (Å) | 57.35 | 50.26 | 47.54 |
| $qR_g$ max | 1.29 | 1.29 | 1.29 |
| $D_{max}$ (Å) | 215 | 190.88 | 181.90 |

| | | | |
|---|---|---|---|
| $\chi^2$ (total estimate from GNOM) | 0.51 | 0.68 | 0.69 |
| Porod volume, Po | 446253 | 278444 | 189214 |
| **R factor analysis** | | | |
| R factor $= \frac{\sum \lvert I(q_i) - I_{model}(q_i) \rvert}{\sum \ I(q_i)}$ (Extended model, coiled-coil) | 1.6 %, 2.5 % | 2.0 %, 2.3 % | 4.3 %, 1.8 % |
| Weighted R factor $\left(\frac{\sum \ (I(q_i) - I_{model}(q_i))^2/\sigma(I(q_i))^2}{\sum \ I(q_i)^2/\sigma(I(q_i))^2}\right)^{0.5}$ (Extended model, coiled-coil) | 3.6 %, 3.4 % | 5.4 %, 4.2 % | 3.6 %, 2.3 % |
| **SASDB accession codes** | SASDJW9 | SASDJX9 | SASDJY9 |

**Table S8: List of primers for MtMce1A-1F constructs**

| **Construct Name** | **Primers** |
|---|---|
| MtMce1A-FP-Nco1 | 5' TGT ACC ATG GCA ACG ACG CCG GGG AAG CTG AAC 3' |
| MtMce1A-RP-Xho1 | 5' CCA GCT CGA GTA ATG GGT TGA TCG TGT TAT CCC CTA CCT G 3' |
| MtMce1B-FP-Nco1 | 5' CAC GCC ATG GCA AAA ATC ACT GGA ACC GTC GTC AAA CTC 3' |
| MtMce1B-RP-Xho1 | 5' TCC ACT CGA GTA ATT GCG GCG TGC ACC TAC CCG 3' |
| MtMce1C-FP-Nco1 | 5' GTG CCC ATG GCA AGA ACG CTG GAA CCA CCC AAC 3’ |
| MtMce1C-RP-Xho1 | 5' AAG AGA GCT CTA AAT TCT CGC TAC CTC CCG TCA CGC C 3' |
| MtMce1D-FP-Nco1 | 5' AGG TCC ATG GCA TTG AGC ACC ATC TTT GAT ATC CGC 3' |
| MtMce1D-RP-Xho1 | 5' CAG CAC TCG AGT AAT TGA CCC CCT CCT GCC TCA G 3' |
| MtMce1E-FP-Nco1 | 5' AGG ACC ATG GCA ATG AGC GTG CTG GCG CGG AT 3' |
| MtMce1E-RP-Hind III | 5' ATG AAA GCT TTA AGC ACT GGC GAT TTC CCC TTT CTA CCA GCG G 3' |
| MtMce1F-FP-Nco1 | 5' CAG CAC TCG AGT AAT TGA CCC CCT CCT GCC TCA G 3' |

| | |
|---|---|
| MtMce1F-RP-Xho1 | 5' TCG GAA GCT TTA AGC TGG CCG GCG CCA GCA TCT C 3' |
| MtMce1A$_{38\text{-}454}$ FP-Nco1 | 5’ ATA CCC CAT GGC ACG CGG GGA GTT CAC 3’ |
| MtMce1A$_{38\text{-}454}$ RP-Xho1 | 5’ CCA GCT CGA GTA ATG GGT TGA TCG TGT TAT CCC CTA CCT G 3’ |
| MtMce1B$_{29\text{-}346}$ FP-Nco1 | 5' GTG ACC ATG GCA CAG ATG CGC TTC GAC CGG AC 3’ |
| MtMce1B$_{29\text{-}346}$ RP-Xho1 | 5' TCC ACT CGA GTA ATT GCG GCG TGC ACC TAC CCG 3' |
| MtMce1D$_{44\text{-}314}$ FP | 5'- CTC TCC ATG GCA CTG ACG AAC AAC ACG GTG GTC GCC -3' |
| MtMce1D$_{44\text{-}314}$ RP | 5' GGT ACT CGA GGT TAA TGT TCG CCG CCA GCG TC 3' |
| MtMce1E$_{37\text{-}390}$ FP | 5’CTACTGAGAATCTTTATTTTCAGGGCGCCATGTCCAATGTGGCGATCCCCGG3’ |
| MtMce1E$_{37\text{-}390}$ RP | 5’ GTGGTGGTGGTGGTGCTCGAGTAAGCACTGGCGATTTCCCC 3’ |
| MtMce1F$_{30\text{-}314}$ FP | 5' CTG GTC CAT GGC ACG AAT TCC GAG TCT GGT GGG TAT CGG GC 3' |
| MtMce1F$_{30\text{-}314}$ RP | 5' GAA TAC TCG AGC GCC GGC GCG AGC GGC -3' |
| MtMce1A$_{36\text{-}148}$ FP | 5’ CTTTATTTTCAGGGCGCCATGGCACAGTTTCGCGGGGAGTTCACG-3’ |
| MtMce1A$_{36\text{-}148}$ RP | 5’ GTGGTGGTGGTGGTGCTCGAGCTCGGTGGTCACCGACCGTACGTCG-3’ |
| MtMce1A$_{126\text{-}454}$ FP | 5’ CTTTATTTTCAGGGCGCCATGGCA CCGAAAAACCCGACAAAGAGGCGG 3’ |
| MtMce1A$_{126\text{-}454}$ RP | 5’ GTGGTGGTGGTGGTGCTCGAG TAA TGG GTT GAT CGT GTT ATC CCC TAC C 3’ |
| MtMce1A$_{38\text{-}325}$ FP | 5’ CTT TAT TTT CAG GGC GCC ATG GCA CGC GGG GAG TTC ACG CCC AAG3’ |
| MtMce1A$_{38\text{-}325}$ RP | 5' GTG GTG GTG GTG GTG CTC GAG TGA GTT CGT CCT CAG CGA GTA GCC G 3' |

**Table S9: List of primers for MtMce4A-4F constructs**

| Construct Name | Primers |
|---|---|
| MtMce4A-FP-Nco1 | 5' CAG ACC ATG GGG TCC GGC GGC GGA TCT CGA C 3' |
| MtMce4A-RP-Xho1 | 5' GAG CCT CGA GTA AGA AGT CGT CCC GTT CCG CGA ACG 3' |
| MtMce4B-FP-Nco1 | 5' CTT CCC ATG GCG GGC TCG GGC GTT CCC 3' |
| MtMce4B-RP-Xho1 | 5' ACT TCT CGA GCA ATT TAG CAA AGG CGC ACC TCC CCT TGC 3' |
| MtMce4C-FP-Nco1 | 5' GGA GCC ATG GCA TTG CTA AAT AGG AAG CCA AGT AGC 3' |
| MtMce4C-RP-Xho1 | 5' ACC CCT CGA GTA ACG GCG ACT TCG GTC TGA 3' |
| MtMce4D-FP-Nco1 | 5' ACC GCC ATG GCA GTG ATG GGC CGG GTC GCC ATG 3' |
| MtMce4D-RP-Xho1 | 5' CAG ACT CGA GTA ATC CGC CGC CCC CAT GCT CGC CCG 3' |
| MtMce4E-FP-Nco1 | 5' TGG GCC ATG GCA AAC CGA ATC TGG TTG CGC GCC 3' |
| MtMce4E-RP-Hind III | 5' GTC CAA GCT TCA ACT GTC CCG ACG CCG TAC CGG GTG 3' |
| MtMce4F-FP-Nco1 | 5' AGG GCC ATG GCA ATG ATC GAC CGA CTC GCC AAG 3' |
| MtMce4F-RP-Xho1 | 5' TCT GCT CGA GCA ACA GCT GCC TCG GAT CGC GC 3' |

| | |
|---|---|
| MtMce4A$_{36-400}$ FP | 5' GTG ACC ATG GCA CAG ATG CGC TTC GAC CGG AC 3' |
| MtMce4A$_{36-400}$ RP | 5' TCC ACT CGA GTA ATT GCG GCG TGC ACC TAC CCG 3' |
| MtMce4A$_{39-140}$ FP | 5' TTATTTTCAGGGCGCCATGGCCTCTACGGACACCGTCACGGTAT 3’ |
| MtMce4A$_{39-140}$ RP | 5' GTGGTGGTGGTGGTGCTCGAG CTC AAG CTG TAC CTG AGA CGC C 3' |
| MtMce4A$_{121-400}$ FP | 5' TATTTTCAGGGCGCCATGGCCAAGACGCCGTCGCCCAAGCCG 3' |
| MtMce4A$_{121-400}$ RP | 5' GTGGTGGTGGTGGTGCTCGAGGAAGTCGTCCCGTTCCGCGAAC 3' |
| MtMce4A$_{39-320}$ FP | 5’ TTATTTTCAGGGCGCCATGGCCTCTACGGACACCGTCACGGTAT 3’ |
| MtMce4A$_{39-320}$ RP | 5' GTGGTGGTGGTGGTGCTCGAG CAA CAC GAA GCT CGA CGA GGT G 3' |
| MtMce4A$_{321-400}$ FP | 5’ AGAATCTTTATTTTCAGGGCGCCATGGCC GGT GCG CCG TCG TAC ACC TAT CC 3’ |
| MtMce4A$_{321-400}$ RP | 5'-GTGGTGGTGGTGGTGCTCGAGGAAGTCGTCCCGTTCCGCGAAC 3' |

**Table S10: List of buffers used in SEC purification**

| Protein Name | Buffer |
|---|---|
| MtMce4A, MtMce4C, MtMce4D | 50 mM Tris, 500 mM NaCl, 10% Glycerol, 5mM DDM, 1 mM β-ME, pH 8.0 |
| MtMce4B, MtMce4E, MtMce4F | 50 mM Tris, 500 mM NaCl, 10% Glycerol, 5 mM FC-12, 1 mM β-ME, pH 8.0 |
| MtMce4A$_{36-400}$, MtMce4A$_{121-400}$, | 50 mM Tris, 500 mM NaCl, 10% Glycerol, 5mM DDM, 1 mM β-ME, pH 8.0 |
| MtMce4A$_{39-320}$ | 50 mM Tris, 500 mM NaCl, 10% Glycerol, 5mM DDM, 1 mM β-ME, pH 8.5 |
| MtMce4A$_{39-140}$ | 50 mM MOPS, 350 mM NaCl, 10% Glycerol, pH 7.0 |
| MtMce1A$_{36-148}$ | 50 mM TRIS, 350 mM NaCl, 10% Glycerol, pH 8.5 |
| MtMce1A$_{38-325}$ | 50 mM Tris, 500 mM NaCl, 10% Glycerol, 5mM DDM, 1 mM β-ME, pH 7.5 |
| MtMce1A$_{126-454}$ | 50 mM Tris, 350 mM NaCl, 10% Glycerol, 5mM DDM, 1 mM β-ME, pH 7.5 |
| MtMce1A$_{38-454}$ | 50 mM Tris, 500 mM NaCl, 10% Glycerol, 5mM DDM, 1 mM β-ME, pH 8.0 |
| MtMce1B$_{29-346}$ | 20 mM Hepes, 500 mM NaCl, 10% Glycerol, 15mM Fos Choline-12, 1 mM β-ME, pH 7.0 |
| MtMce1C, MtMce1D$_{44-314}$ | 50 mM Tris, 300 mM NaCl, 10% Glycerol, 5mM DDM, 1 mM β-ME, pH 8.0 |
| MtMce1E$_{37-390}$* | 50mM Tris, 500mM NaCl, 0.05% C12E9, 1 mM β-ME, pH 8.0 |
| MtMce1F$_{30-314}$ | 50 mM Tris, 500 mM NaCl, 10% Glycerol, 5mM DDM, 1 mM β-ME, pH 8.0 |
| Lysis buffers for all the purified constructs in DDM and FC-12 had 25 mM of the respective detergents, along with protease inhibitor, lysozyme, DNase and RNase. *The lysis buffer had 25 mM DDM in this case. | |
